# Domain coupling in activation of a family C GPCR

**DOI:** 10.1101/2024.02.28.582567

**Authors:** Naomi R. Latorraca, Sam Sabaat, Chris Habrian, Julia Bleier, Cherise Stanley, Susan Marqusee, Ehud Y. Isacoff

## Abstract

The G protein–coupled metabotropic glutamate receptors form homodimers and heterodimers with highly diverse responses to glutamate and varying physiological function. The molecular basis for this diversity remains poorly delineated. We employ molecular dynamics, single-molecule spectroscopy, and hydrogen–deuterium exchange to dissect the pathway of activation triggered by glutamate. We find that activation entails multiple loosely coupled steps and identify a novel pre-active intermediate whose transition to the active state forms dimer interactions that set signaling efficacy. Such subunit interactions generate functional diversity that differs across homodimers and heterodimers. The agonist-bound receptor is remarkably dynamic, with low occupancy of G protein–coupling conformations, providing considerable headroom for modulation of the landscape by allosteric ligands. Sites of sequence diversity within the dimerization interface and diverse coupling between activation rearrangements may contribute to precise decoding of glutamate signals and transients over broad spatial and temporal scales.

## Main

Two classes of neuronal receptors––the metabotropic and ionotropic glutamate receptors (mGluRs and iGluRs)––mediate synaptic transmission and synaptic plasticity (*1*). These multi-domain proteins assemble into homo- and heteromeric complexes of multiple receptor subtypes, with distinct ligand affinity, efficacy, and kinetics that tailor responses to glutamate, within and near the synapse (*2*, *3*). Here, we investigate the molecular basis for allostery in the dimeric mGluR G protein–coupled receptors (GPCRs). Each subunit in the mGluR dimer possesses an extracellular ligand binding domain (LBD) with a clamshell-like topology that links to a seven-pass transmembrane domain (TMD) via a cysteine-rich domain (CRD). Cryo-electron microscopy of full-length mGluRs has identified two distinct conformations, revealing the major rearrangements that occur upon ligand binding: the clamshell of each subunit closes on ligand, and the two clamshells twist relative to the other about the upper LBD; these motions bring the lower LBD surfaces and CRDs into contact to alter packing of the second extracellular loop in the TMD, an effect that enables G-protein coupling at the intracellular surface (*4–9*).

In addition to these two major conformations, recent structural studies reveal a surprising degree of conformational diversity across mGluRs, including an asymmetric arrangement in which only one subunit is closed around ligand and multiple antagonist-bound conformations that differ in the packing and orientation of transmembrane helices (*6*, *10*). Crystal structures of isolated LBD dimers reveal additional conformations, including arrangements in which the two subunits have twisted relative to one another despite both clamshells remaining open, and those in which clamshells have closed in the absence of intersubunit twisting (*4*, *11*, *12*). Similarly, single-molecule spectroscopy studies indicate the existence of three or more extracellular domain conformations, heterogeneity in the proximity of CRDs and TMDs, and decoupling between upper- and lower-lobe LBD motions (*13–20*). Whether and how each of these discrete domain rearrangements––ligand-binding domain closure, intersubunit twisting within the ligand-binding domain, and rearrangement of the cysteine rich domain linkers––couple to one another, and the influence of agonist binding on these equilibria, has yet to be determined.

Varied responses of mGluR homo- and hetero-dimers to glutamate suggest that subunit interactions shape the conformational landscape to tune activation. In group I and II homodimers, glutamate binding to only one subunit greatly reduces the degree and speed of activation, indicating that activation is positively cooperative (*13*, *21*, *22*). In Group III homodimers, glutamate fails to fully stabilize the active conformation, but the group II/III mGluR2/7 heterodimer has accelerated activation kinetics and reaches full efficacy even when liganded at only one of its subunits (*13*, *23*). We recently demonstrated that this conformational diversity arises, at least in part, from variation in low sequence-conservation regions at subunit interfaces (*24*), but how this diversity influences conformational coupling across the various mGluR homo- and heterodimers remains unknown. We combine single-molecule Förster resonance energy transfer (smFRET), molecular dynamics (MD) simulations, and hydrogen–deuterium exchange monitored by mass spectrometry (HDX-MS) to determine the atomic-level mechanisms by which ligand activates the receptor.

We find that three extracellular rearrangements associated with global activation––closure of individual ligand-binding-domain (LBD) subunits, twisting of the LBD subunits relative to one another, and approach of the cysteine-rich domain (CRD) linkers––occur in a loosely coupled manner, such that these events need not occur concomitantly. Our single-molecule studies reveal the existence of a clamshell-closed intermediate that lacks LBD twisting; additionally, in the agonist-bound, LBD-twisted state, the receptor remains dynamic, sampling multiple distinct states associated with varying degrees of proximity of the CRDs. Thus, the receptor populates a G protein–coupling conformation only a fraction of the time. Simulations similarly capture the existence of a pre-active, clamshell-closed conformation of the LBD dimer and reveal key interface interactions whose formation favor the twisted, proximal orientation of the two LBDs. Complex changes in amide hydrogen exchange in the presence of ligand reflect direct stabilization of helices connected to the ligand binding pocket. This observation suggests a possible mechanism by which closure of one subunit favors closure and reorientation of the partner subunit. Moreover, mGluR heterodimers exhibit distinct transition kinetics associated with clamshell closure, providing a mechanistic explanation for the wide variation of signaling efficacies observed across different mGluR heterodimers.

## Results

### Ligand binding domain closure is incompletely coupled to LBD reorientation

To study agonist-induced rearrangements in the LBD, we carried out smFRET measurements in mGluR2 using amber codon suppression to incorporate unnatural amino acids (UAAs) for dye addition via click chemistry (Fig. 1A). To monitor clamshell closure, we incorporated amber stop codons into two sites within one LBD subunit, which increase in proximity when that subunit closes around ligand: Ser463 on the upper lobe and Gln359 on the lower lobe. We co-transfected this construct with an unaltered mGluR2 construct containing an N-terminal HA tag, along with a tRNA/tRNA synthetase pair and the UAA TCO-lysine (see Methods). Receptors were isolated following concurrent on-cell labeling with tetrazine dyes and biotinylated anti-HA antibody. This strategy enabled us to isolate cell surface–expressed mGluR2 with one doubly labeled and one unlabeled subunit via immunoprecipitation of the unlabeled subunit and, thus, exclusively monitor the conformational behavior of the labeled subunit. To monitor LBD twisting, we incorporated an amber stop codon at Ala248 and monitored the distance between the same site on the lower lobe of the two subunits. We note that a recent study employed nearly identical reporter sites to monitor upper and lower-lobe closure and twisting on a sub-millisecond time-scale (*20*); here, we obtain FRET trajectories for single molecules over seconds, enabling us to directly observe conformational transitions between stable states in both mGluR homo- and heterodimers. Below, we integrate observations from smFRET with higher-resolution biophysical approaches to reveal structural features that couple LBD closure and twisting to set ligand efficacy.

**Figure 1.**
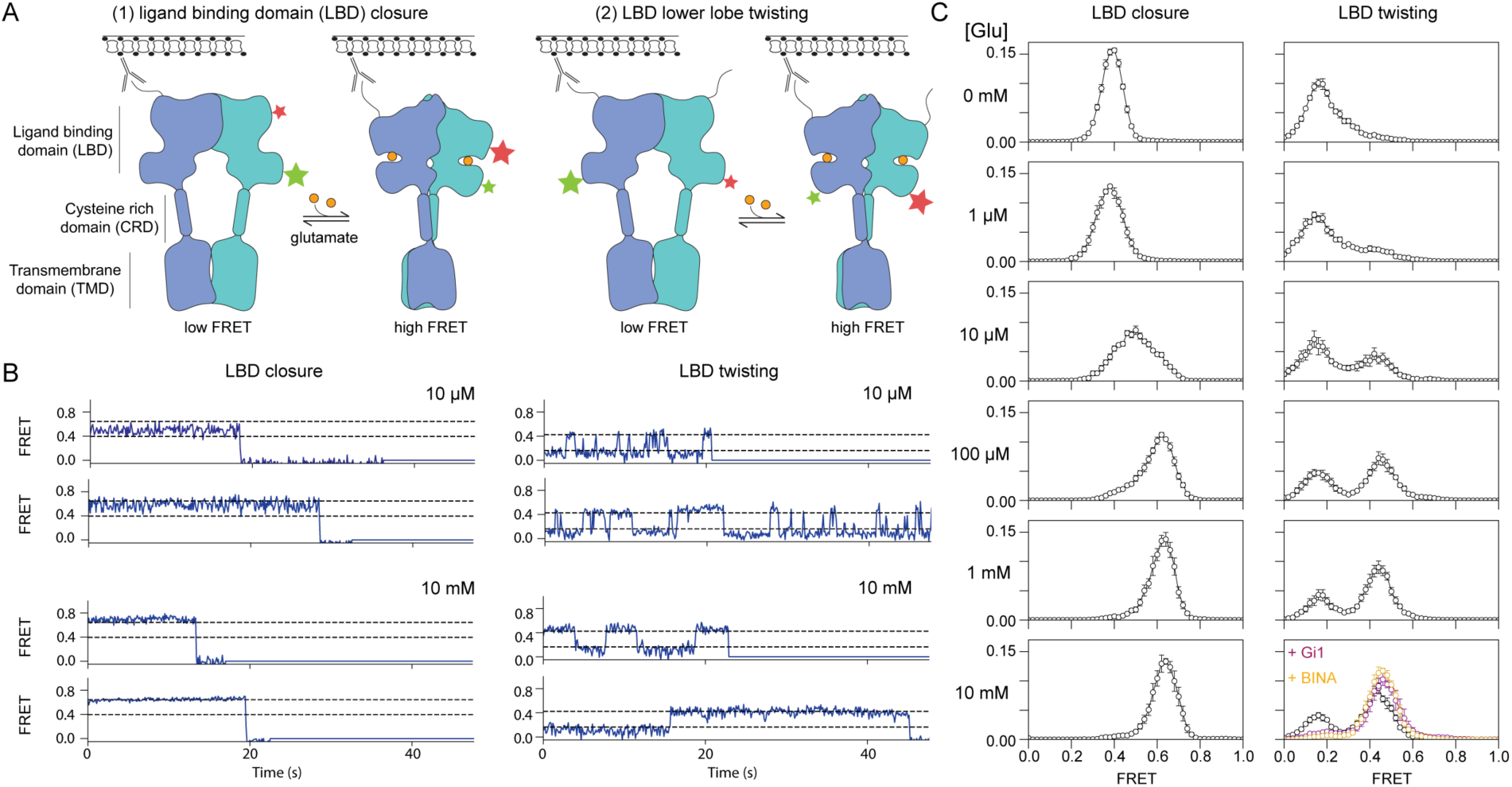
Agonist-induced LBD closure is loosely coupled to intersubunit activation of the extracellular domain. (A) Donor/acceptor pairs used to distinguish between intrasubunit LBD closure (left, middle-left) and intersubunit twisting (middle-right, right). For both pairs, adding agonist brings donor/acceptor pairs into closer proximity, thereby increasing the FRET signal. Detergent-solubilized receptors undergo stochastic labeling with donor and acceptor fluorophores via click chemistry, followed by low-density immobilization on a coverslip and single-molecule TIRF imaging. Analysis is limited to puncta with a single donor and single acceptor. (B) single-molecule FRET traces for the two FRET pairs, carried out in the presence of EC50 (10 µM) and saturating (10 mM) levels of glutamate. (C) smFRET histograms collected under a range of glutamate concentrations for each FRET pair. Lower right: 10 mM glutamate alone (black symbols) and along with either 100 µM of the positive allosteric modulator BINA (orange) or 4 µM of Gi1 (red). ≥ 4 movies per histogram; error bars represent standard error of the mean (s.e.m.).

We monitored clamshell closure and LBD lower-lobe twisting across a range of glutamate concentrations (Fig. 1C, left; Extended Data Fig. 1). At 0 mM Glu, the clamshell sensor had a narrow symmetric distribution that populated a low-FRET state (E ∼ 0.38), with single receptor traces showing a stable FRET level, consistent with a single, open conformation. At 10 mM Glu, the clamshell sensor populated a high-FRET state (E ∼ 0.63), consistent with a single, closed conformation. At an intermediate glutamate concentration of 10 µM Glu, the FRET distribution broadened symmetrically centered around an intermediate FRET peak (E ∼ 0.50), due to rapid interconversion between open and closed conformations, consistent with this concentration being close to the EC50 of mGluR2 (*25*).

The LBD twisting sensor revealed a distinct set of occupancies in response to increasing glutamate concentration (Fig. 1C, right). In the absence of glutamate, this sensor primarily populated a low-FRET state (E ∼ 0.18), but the distribution was broad and skewed to higher FRET values. Addition of glutamate resulted in occupancy of a second, high-FRET state (E ∼ 0.44). However, full occupancy of the high-FRET twisted state was not achieved even at 10mM Glu, a concentration at which the clamshell sensor was fully closed (Fig. 1B, C). Instead, individual trajectories switched between the low-FRET, relaxed conformation and high-FRET active conformation, with long-lived dwells in the high-FRET state (τ ∼ 3.0 s, where τ is the inverse of the average number of transitions per second; see Methods) (Fig. 1B; Extended Data Fig. 1). Thus, although glutamate binding favors both clamshell closure and intersubunit twisting, these two conformational rearrangements are not coupled through a single rigid-body movement.

These observations indicate that clamshell closure is *not* sufficient to fully stabilize the activated, twisted orientation, leaving the LBDs to transition between two conformations, both of which possess closed LBDs (C/C): an intermediate conformation, with LBDs in a relaxed orientation (R-C/C), and an activated conformation, with LBDs twisted towards one another (A-C/C). Here, we use “relaxed” or (R) to refer to any conformation in which the distance between the two lower lobes matches the distance observed in inactive-state (R-O/O) structures. Notably, addition of either the positive allosteric modulator (PAM) BINA, which binds within the transmembrane domain (TMD), or of the Gi heterotrimer, which binds the TMD intracellular surface, stabilizes the high-FRET twisted conformation of the LBD, consistent with assignment of this conformation to the active state (Fig. 1C) (*26*, *27*). To further probe the activation mechanism, we next sought to identify the structural determinants that regulate the transition to this fully activated LBD conformation.

### Dimerization interface interactions set mGluR efficacy

In ligand-bound, clamshell-closed states, what structural features maintain the ‘twisted’, active orientation, observed in agonist-bound cryo-EM structures? We reasoned that all-atom, molecular dynamics simulations initiated from the active-state structure would (1) reveal persistent residue– residue interactions that favor the active, twisted state, and (2) transition on long-timescales to distinct conformational states, thereby revealing atomic-level mechanisms that, in reverse, correspond to conformational pathway(s) to the active conformation (*28*). To reduce computational cost, we initiated all-atom molecular dynamics simulations of an mGluR2 LBD dimer, which lacks the TMD and CRD, starting from a ligand-bound active-closed/closed (A-C/C) crystal structure (PDB entry 4XAQ) (*29*). We retained the co-crystallized high-affinity, cyclically constrained glutamate analog, LY354740, in each binding pocket, along with co-crystallized monovalent cations and anions and resolved water molecules. In each of five long-timescale simulations (∼15 µs per simulation), the mGluR2 LBD persisted in its initial A-C/C conformation for at least several microseconds. In three of these five simulations, we observed transitions out of the A-C/C conformation to a closed/closed conformation with the lower lobes twisted away from each other after residing for several microseconds in the initial A-C/C conformation (Fig. 2A). In this new conformation, the lower lobes are separated from each other such that the distance between their centers of mass is ∼57 Å, approximately the same separation distance as observed in inactive-state (R-O/O) structures (∼60 Å) (*4–6*) and much greater than the distance (∼34 Å) observed in active-state, twisted (A-C/C) structures. As both clamshells remain closed around ligand, with the lower lobes separated at distances reminiscent of the inactive, R-O/O state, we refer to this new conformational intermediate as a relaxed, closed/closed, or R-C/C, conformation. In this state, the upper lobes form a new dimerization interface in which the plane formed by the B and C helices of one protomer is partially rotated with respect to the plane formed by the B and C helices of the opposite protomer (Fig. 2A). Within the upper-lobe dimerization interface, the number of van der Waals contacts formed between helices B and C of opposing subunits is fewer than those observed in either inactive-state or active-state structures (Extended Data Fig. 7B). The relaxation associated with the transition to an R-C/C intermediate largely occurs through a rigid-body swinging motion: the root-mean-square deviation (r.m.s.d.) of either the upper (residues 22-180, 321-436) or lower (residues 188-320, 463-487) lobe to their initial conformations is less than 1.0 Å, suggesting that the transition occurs with only minor shifts in protein backbone conformation.

**Figure 2.**
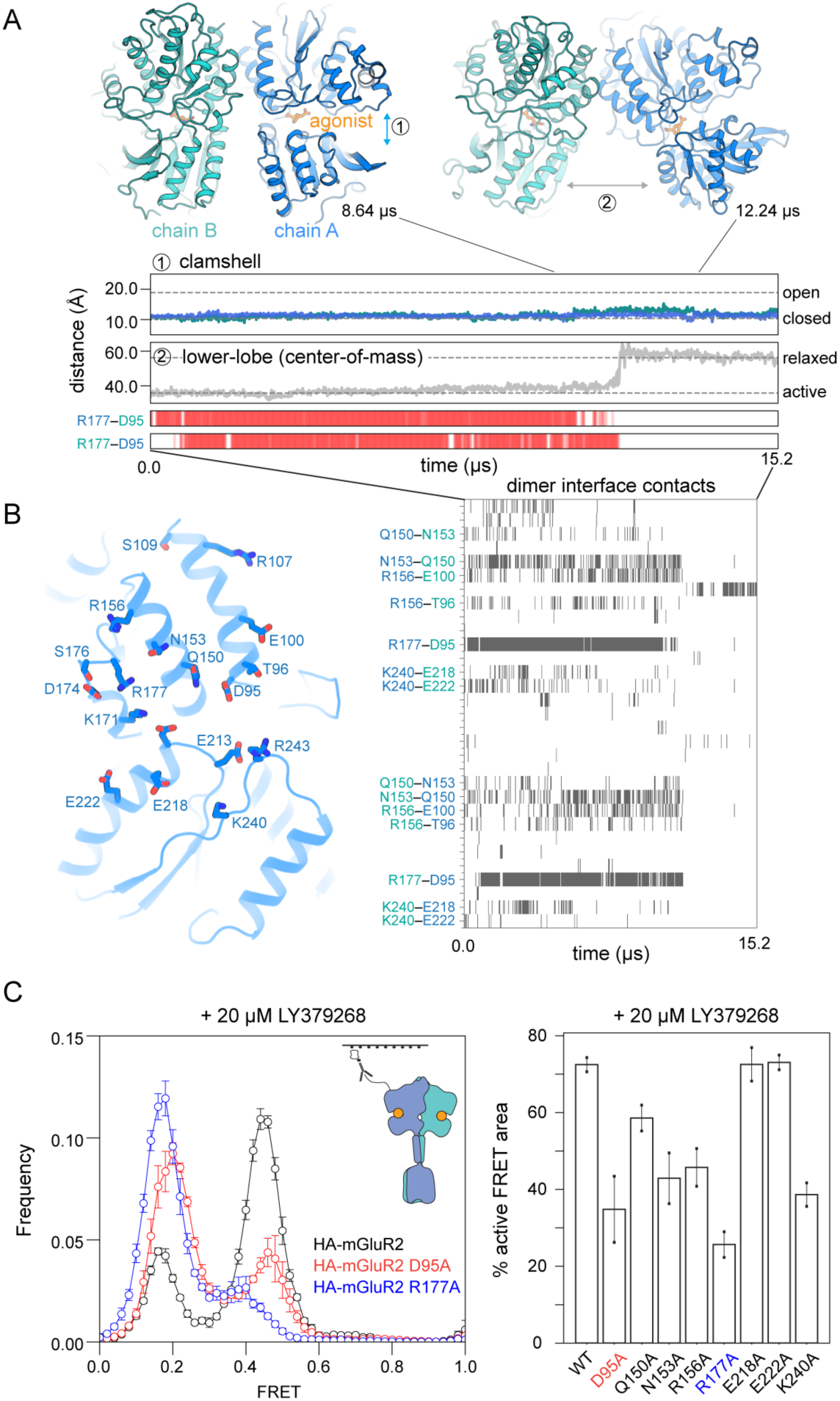
An electrostatic network controls the relaxed–active conformational transition. (A) Snapshots from an MD simulation, before and after the transition from an active, closed conformation to a relaxed, closed conformation. The distance between the upper and lower lobes of the LBD (or ‘clamshell’) is shown for each subunit and remains the same over the course of the simulation; by contrast, the distance between the lower lobe of each subunit increases substantially after the transition to values seen in relaxed-state structures. (B) Residues involved in cross-protomer interactions at the dimerization interface are shown on the structure of the mGluR2 LBD (left), and dimer interface cross-protomer contacts are shown over time using gray bars (downsampled every 12 ns; right). (C) smFRET experiments to monitor inter-subunit twisting reveal that R177A and D95A mutations reduce population of the high-FRET peak (left). Percentage of high-FRET population in smFRET measurements of inter-subunit twisting on additional polar residues at the dimer interface (right); error bars represent standard error of the mean (s.e.m.) across ≥ 5 movies.

To identify the molecular interactions that govern transitions between R-C/C and A-C/C conformations, we examined polar interactions within the dimerization interface that disappeared upon the A-C/C to R-C/C transition. Only a small number of residues formed persistent cross-protomer interactions during active-state portions of simulation, including a network of four residues immediately beneath the hydrophobic interface on the upper lobe surface (E100, R156, Q150, N153); an electrostatic network involving loops at the base of the upper lobe (D95, R177, R243); and an electrostatic network between the lower lobe interfaces (K240, E218, E222) (Fig. 2B). Of these interactions, the two R177–D95 salt bridges were highly stable; disruption of the two R177–D95 pairs was tightly coupled to the transition to the intermediate state. We hypothesized that interactions between these two residues, as well as with R243, which forms an arginine π stack with R177 in one protomer of certain crystal structures, maintain the twisted orientation of the upper-lobe interface in the active state, such that the hydrophobic residues on either side of this interface continue to pack against one another.

We tested the predictions from these simulations in smFRET on alanine variants at key interface residues, including R177 and D95. We found that, in the presence of a saturating amount of the high-affinity agonist LY379268 (a high-affinity analog of LY354740), mGluR2-R177A exhibited a ∼75% decrease in occupancy of the active conformation relative to wild-type mGluR2, while D95A had a similar, albeit smaller, effect (Fig. 2C). Other single alanine mutations to simulation-identified interface residues also had significant effects on the active conformation, although these effects were also smaller than R177A. We also confirmed that, in the presence of LY379268, the individual R177A and D95A variants fully occupied the closed clamshell conformation (Fig. 3A, Extended Data Fig. 3), indicating that the depletion of the active-state population is not due to a loss in agonist-induced clamshell closure. Thus, smFRET mutational analysis provides experimental support for the MD simulations and, together, these demonstrate that a cross-subunit electrostatic interaction network plays a key role in stabilizing the twisted orientation of the closed LBDs, thereby regulating agonist efficacy.

**Figure 3.**
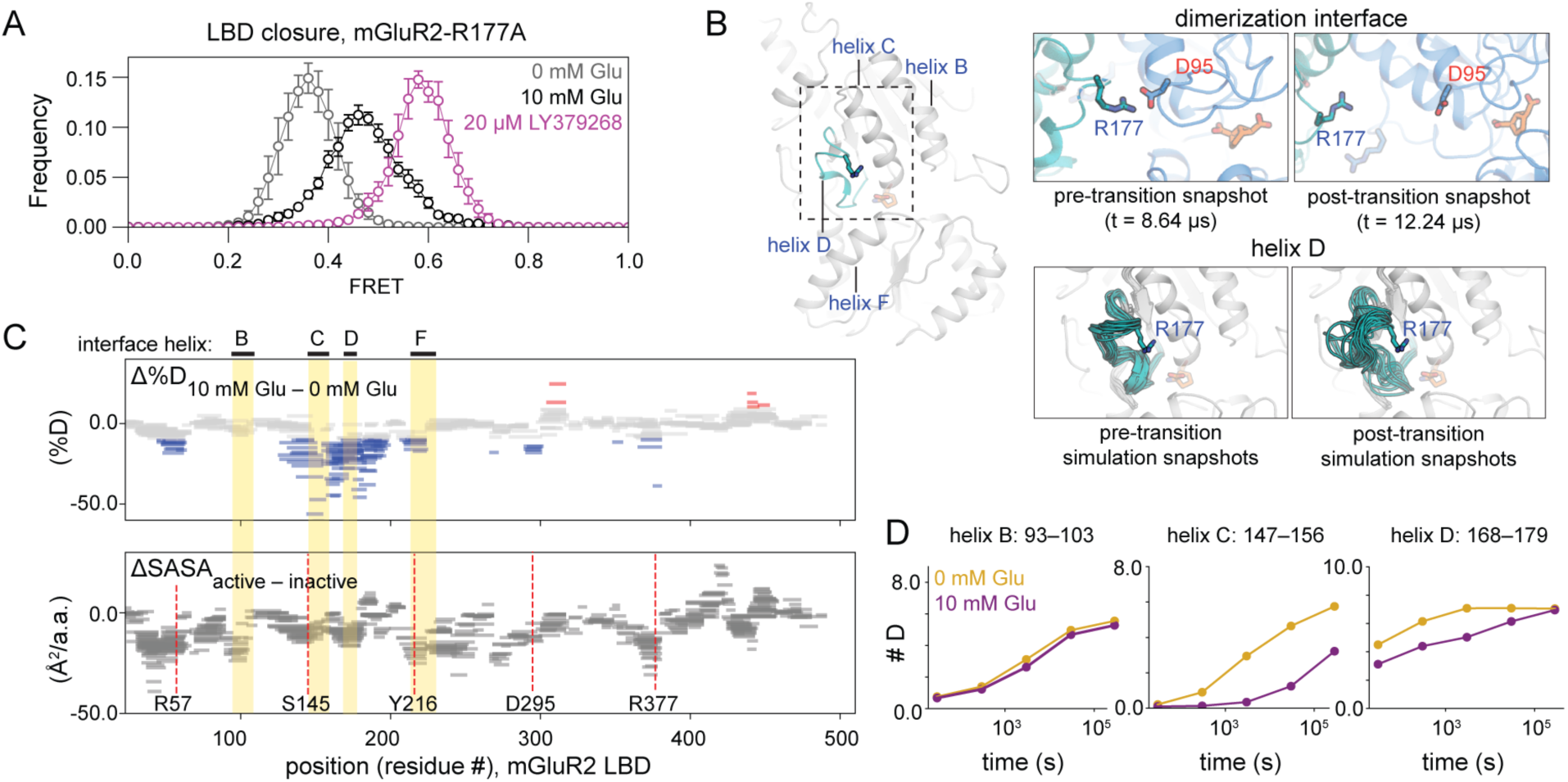
Glutamate binding differentially stabilizes interface helices. (A) A dimerization interface mutant, mGluR2-R177A, exhibits reduced clamshell closure in response to 10 mM glutamate (black) compared to wild-type (compare to Figure 1C, lower left) or to mGluR2-R177A in the presence of high-affinity agonist LY379268 (purple). (B) Snapshots from MD simulation with just the R177-containing helix (helix D) shown (full structure on left) demonstrate how a key salt bridge breaks apart upon transition to the relaxed, closed intermediate (top) and how, after this transition, helix D becomes less ordered (bottom). 10 frames per image, downsampled every 360 ns, before and after the transition. (C) (Top) Woods plots from HDX-MS experiment at 30,000 s: each horizontal bar corresponds to a peptide representing a fragment of the mGluR2 LBD sequence identified by mass spectrometry; the height of each bar corresponds to the peptide’s change in % deuteration in the presence vs. in the absence of 10 mM glutamate, such that negative values (blue) correspond to increased protection in 10 mM glutamate and positive values (red) correspond to decreased protection in 10 mM glutamate. (Bottom) The change in solvent-accessible surface area in an active-state crystal structure (PDB 4XAQ) vs. in a relaxed-state crystal structure (PDB 5KZN), normalized by peptide length (negative values indicate less solvent exposure in active state). Vertical, dashed red lines indicate ligand-contacting residues in the 4XAQ crystal structure. (D) Uptake plots from HDX-MS experiments demonstrate non-uniform changes in protection across interface helices B-D in the presence of 10 mM glutamate (purple) vs. 0 mM glutamate (yellow).

Previous studies indicate that mGluR activation is cooperative, such that ligand binding to both subunits results in more than twice the response of ligand binding to a single subunit (*13*, *21*, *22*, *25*). We therefore sought to determine whether the dimerization interface is responsive to the conformation of the ligand binding pocket (i.e., whether either clamshell is open or closed). Three pieces of evidence suggest a mechanism by which certain helices within the dimerization interface couple intrasubunit conformational changes to intersubunit rearrangements.

First, we found that, unlike the high-affinity agonist LY379268, glutamate alone (up to 10 mM) is not sufficient to fully stabilize the closed clamshell conformation of mGluR2–R177A (Fig. 3A). Thus, although mGluR2-R177A *can* adopt a closed clamshell conformation, and does so readily in the presence of LY379268, the fact that mutation of a residue within the dimerization interface alters clamshell closure suggests that the binding pocket and dimerization interface are conformationally coupled.

Second, in simulation, we found that the short helical loop spanning residues 169–177 (helix D) increases in conformational flexibility after transitioning from the A-C/C conformation to the R-C/C intermediate (Fig. 3B; Extended Data Fig. 2B). Thus, contacts at the dimerization interface that are formed only in the active, twisted orientation aid in stabilizing an ordered conformation of the R177-containing helix.

Third, we hypothesized that additional structural elements in the dimerization interface may also be sensitive to ligand binding, but that fluctuations in those elements may occur on timescales not accessible to MD simulations. We therefore sought to employ an orthogonal experimental method to monitor local conformational flexibility that is not restricted by the timescales of the simulations. We turned to continuous labeling hydrogen–deuterium exchange monitored by mass spectrometry (HDX-MS). HDX-MS reports on the flexibility of regions of the protein via amide protons as they fluctuate from a ‘closed’ (usually, hydrogen-bonded) structure, which is not accessible to solvent, to an ‘open’ conformation, which is accessible for exchange with solvent deuterons. Under most experimental conditions, HDX occurs in what is referred to as an ‘EX2’ regime, in which the observed rate of hydrogen exchange is related to the equilibrium populations of open (exchangeable) and closed (unexchangeable) conformations. In this scenario, slower exchange (also referred to as increased protection) indicates less flexibility or sampling of the open conformation (*30*). Detection of HDX by mass spectrometry allows hydrogen exchange to be measured at the level of individual peptides, giving information on local flexibility. We carried out HDX-MS on the mGluR2 LBD, in the presence and absence of glutamate, to enable comparisons to our LBD-only simulations.

Several regions of the mGluR2 LBD exhibit slowed exchange (increased protection) in the presence of glutamate compared to in its absence (Fig. 3B,C, Extended Data Fig. 4). For example, peptides that contain residues that directly contact the ligand show slowed exchange in the presence of glutamate compared to in its absence (Fig. 3B). This is also true for most peptides in the dimerization interface including those in helices C, D and F. However, for helix D peptides containing R177, several protons exchange very quickly, consistent with the increased flexibility seen in MD simulations (Fig. 3B, C). The neighboring helix, helix C (residues 145–158), is in general much less dynamic, with no exchange observed in the first timepoints in the presence and absence of glutamate. Thus, the HDX-MS data suggest that although glutamate stabilizes helix C, this region is already stable even in the absence of glutamate, thereby explaining the absence of notable fluctuations for this region in simulations.

As described above, protection from exchange arises from stabilization of individual hydrogen bonds and/or burial from solvent. In an effort to deconvolve these effects, we compared the observed changes in HDX to the changes in average solvent accessible surface area derived from crystal structures of the two conformations. Structural analysis reveals that all the dimerization interface helices (helices B, C, D and F) show increased burial in the active state, albeit to different extents. However, the observed changes in the HDX profiles do not follow thes same patterns (e.g. Helix B shows the greatest change in average solvent accessible surface area with no notable change in hydrogen exchange, whereas helices C and D show modest changes in solvent accessible surface area with relatively large changes in hydrogen exchange). This lack of correlation between the changes in buried surface area and the changes in hydrogen exchange propensity indicate that ligand binding strongly alters the conformational flexibility of a subset of structural elements within the dimerization interface. We therefore propose that the stabilization of helices C and D by glutamate ensures proper positioning of polar and charged residues involved in the R177 network (including R177, as well as Q150, N153 and R156) to form favorable contacts across the dimerization interface.

### Dynamic CRDs result in loose coupling to active state

We next sought to determine how conformational changes in the LBD couple to conformational changes in the cysteine-rich domains (CRDs), which link the LBD to the TMD, and are positioned apart in resting-state structures but close to one another in active-state, full-length mGluR structures. We incorporated an amber stop codon at Ala548, midway along the two CRD linkers, and used our single-molecule assay to monitor the intersubunit distance between these sites (Fig. 4A, cartoon inset). In the absence of glutamate, the CRD sensor populates a low-FRET state (E ∼ 0.28) (Fig. 4A). In glutamate, the CRD sensor populates a second, medium-FRET state (E ∼ 0.48) whose occupancy increases with glutamate concentration but does not persistently reside in that state even at saturating glutamate (10 mM). Instead, it exhibits a bimodal FRET distribution (Fig. 4A), as seen in the lower-lobe twisting sensor (Fig. 1C).

**Figure 4.**
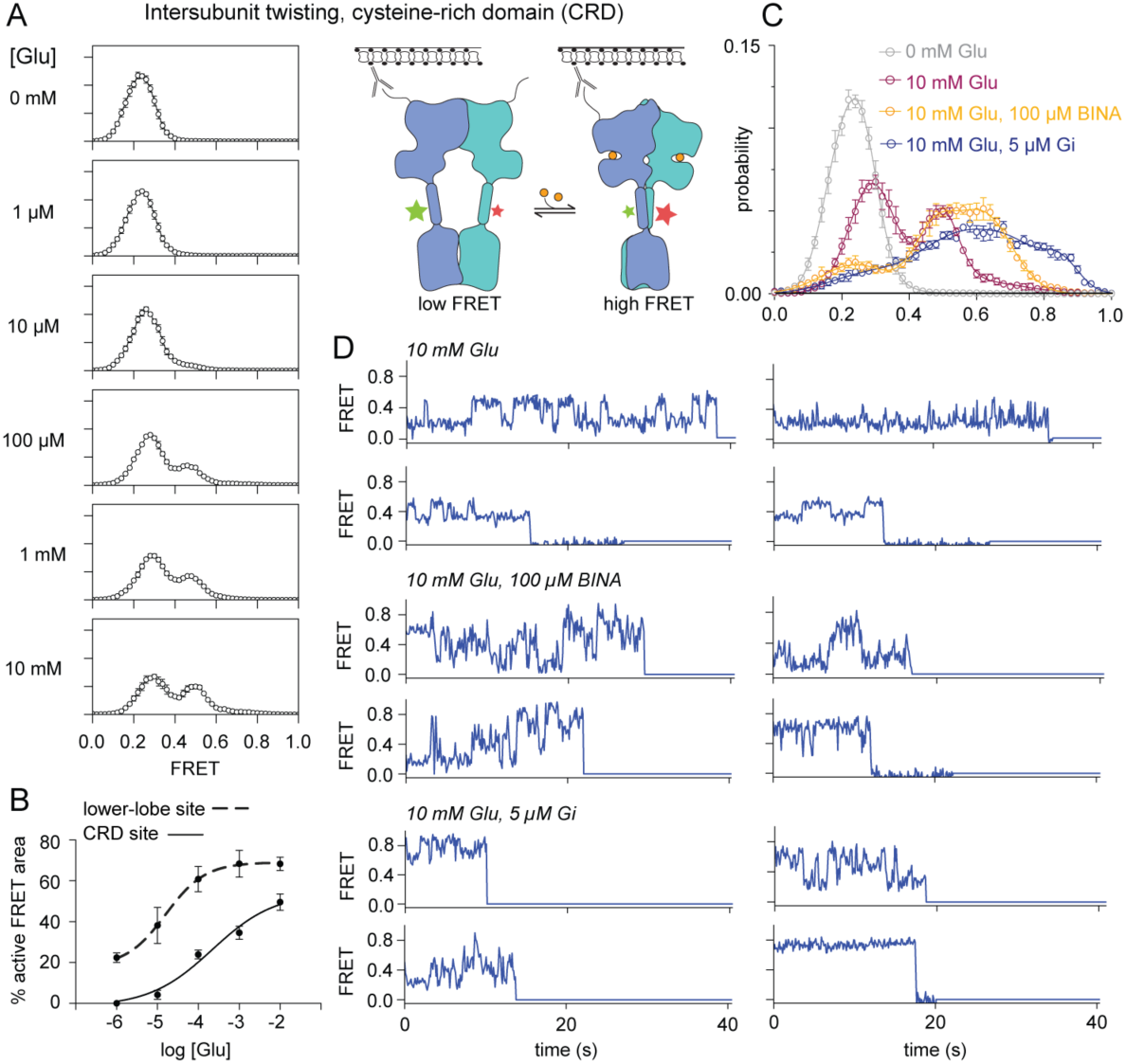
CRD linker acts as a brake on activation. (A) smFRET histograms across increasing concentrations of glutamate, as in Figure 1, using the CRD twisting sensor. (B) Proportion of each smFRET distribution occupying the higher of two FRET states, for either the lower lobe reporter, shown in Fig. 1 (dashed line), or the CRD linker (solid line); bimodal fit carried out for on histogram generated from each movie, with error bars corresponding to S.E.M. for 4–5 movies per concentration. (C) Addition to 10 mM glutamate of the positive allosteric modulator BINA (orange) introduces a higher FRET peak at ∼0.65, whereas addition of the inhibitory heterotrimeric G protein (Gi, blue) spreads to even higher FRET values. 0 mM glutamate (grey) and 10 mM glutamate (purple) replot smFRET distributions from panel (A). (D) Four representative smFRET traces for each condition shown in panel (C).

Despite this similarity, CRD behavior deviates from LBD behavior in four ways. First, while at 10 µM glutamate the lower-lobe LBD sensor has substantial occupancy in the high-FRET state (Fig. 1C), the CRD sensor remains almost entirely in a low-FRET state (E ∼ 0.28) (Fig. 4A, B). Second, while the CRDs approach each other as glutamate concentration increases, leading to population of a prominent medium FRET state (E ∼ 0.48) (Fig. 4A), the dwell time in this state is shorter than the LBD twist high-FRET dwell time (Extended Data Fig. 5C). Third, at 10 mM glutamate, the fractional occupancy of the CRD medium FRET state is ∼45% (Fig. 4A, B), considerably lower than the ∼70% occupancy of the lower-lobe LBD sensor high-FRET state (Fig. 1C, right; Fig. 4B). Fourth, in 10 mM glutamate, the CRDs populate a third conformation of even greater proximity that yields a minor ‘medium-high’ FRET state (E ∼0.65) (Fig. 4C) that can be seen in rare single molecule excursions (Extended Data Fig. 5A). These observations suggest that the LBD and CRD are loosely coupled, and that the receptor can dwell in a conformation with clamshells closed and lower LBD lobes twisted towards each other but with the CRDs spread apart.

Occupancy of the CRD high-FRET state (E ∼ 0.65) is greatly increased at the expense of the low-FRET state (E ∼ 0.28) by addition of a PAM (BINA) to 10 mM glutamate (Fig. 4C, D). Addition of the Gi1 heterotrimer to 10 mM glutamate also increased occupancy of the medium-high FRET state (E ∼ 0.65) and, moreover, favored excursions to an even higher FRET state (E ∼ 0.85) (Fig. 4C, D). Single-molecule traces demonstrate that this highest-FRET active conformation was occupied from hundreds of milliseconds to seconds but that receptors constantly transitioned between this state and all other CRD conformations, resulting in ∼ 20% occupancy of this state (Fig. 4C, D). By contrast, the lower LBD sensor showed almost complete occupancy of the high-FRET twisted state in both BINA and Gi (Fig. 1D), suggesting that all but the low-FRET state of the CRD allosterically stabilizes the LBD twisted state. We interpret this highest-FRET state (E ∼ 0.85) to correspond to the G protein–signaling state. Indeed, in Gi-bound cryo-EM structures, Cα– Cα distances between the corresponding labeling sites span the range of 34–43 Å; a FRET efficiency of E ∼ 0.85, with an R_0_ of 5.1 nm, results in a dye-pair distance of 38 Å, well within this range (Extended Data Fig. 5B).

### Cooperative effects of subtype-specific heterodimers

mGluR2 heterodimerizes with Group II and Group III mGluRs, giving rise to heterodimers with different dimerization propensities, ligand sensitivity and activation kinetics (*13*, *23*, *24*, *31*). For example, using a fusion protein–based FRET sensor attached to either N-terminus that reports on intersubunit twisting, previously we found that glutamate only partially stabilizes the activated conformations of mGluR4/4 and mGluR7/7 but completely stabilizes the activated conformation of mGluR2/7, even with agonist bound to only one subunit (*13*). These data raise the possibility that the conformation of an mGluR subunit may undergo differential modulation through pairing with distinct mGluR subtypes. To test this prediction, we compared mGluR2 clamshell closure energetics in heterodimers containing either mGluR3, mGluR4 or mGluR7 (Fig. 5A). In intermediate glutamate (10 µM), mGluR2 closure kinetics differed between the heterodimers, such that mGluR2/3 and mGluR2/4 displayed longer dwell times in both low- and high-FRET states, whereas mGluR2/2 and mGluR2/7 fluctuated rapidly between states (Fig. 5B, Extended Data Fig. 6). Moreover, occupancy of the clamshell-closed conformation of mGluR2 depended on the partner subunit, with mGluR2 closed a greater fraction of the time when its partner was mGluR3 than when it was mGluR4 (Fig. 5A), consistent with our previous observation that the active conformation of mGluR3 is more stable than that of mGluR2 (*25*).

**Figure 5.**
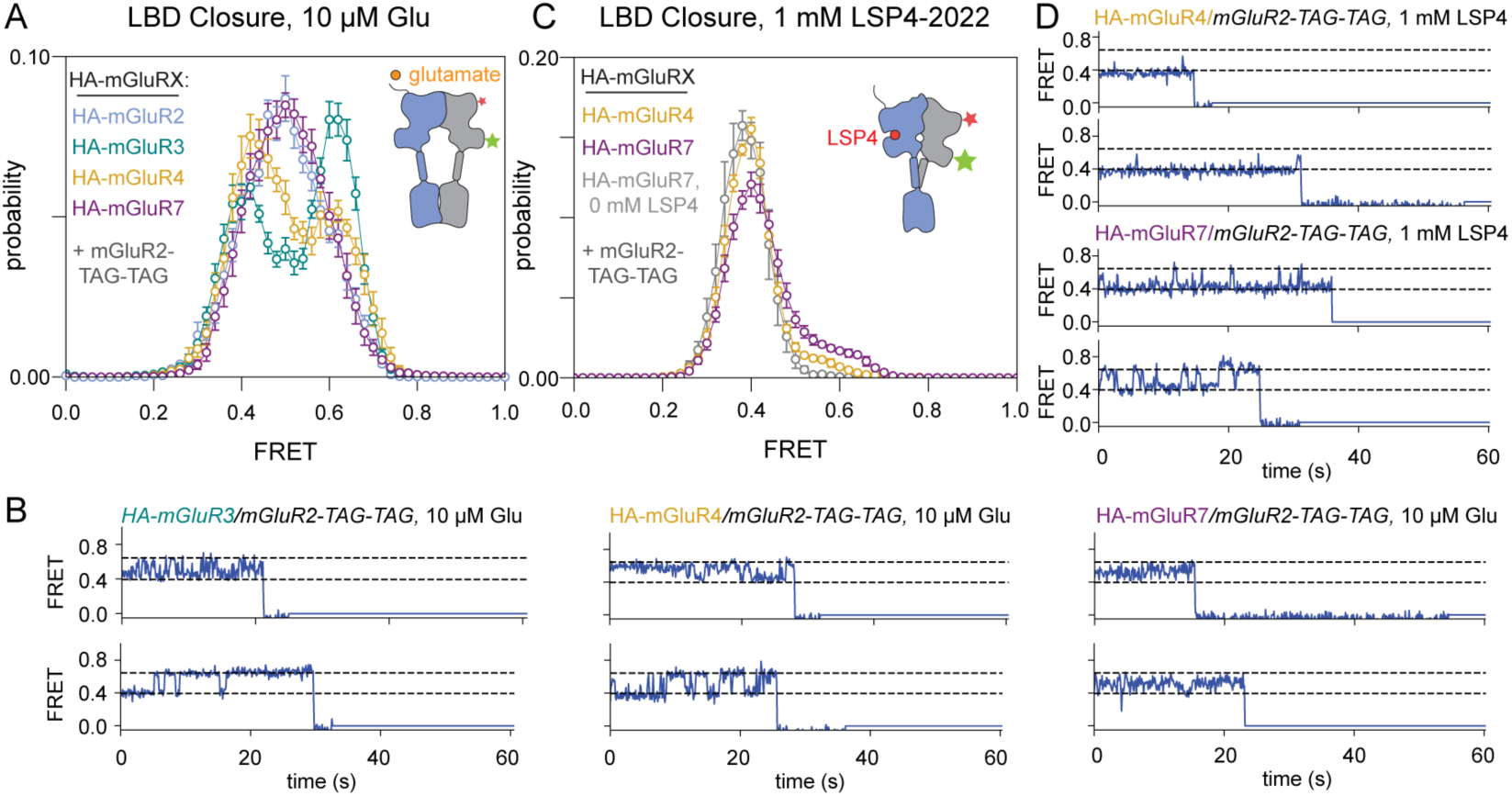
Differential population of mGluR2 LBD-closed states across mGluR heterodimers. (A) FRET distributions in 10 µM clearly resolve open (low-FRET) and closed (high-FRET) peaks only in mGluR2/3 and mGluR2/4 heterodimers. (B) Representative traces from datasets in (A) show distinct mGluR2 LBD open–closed kinetics with different partner subunits. mGluR2/3 and mGluR2/4 display the longest-lived dwells in both LBD open and closed states, in agreement with the well-resolved FRET peaks in (B). (C) In the presence of Group III–specific agonist LSP4-2022, ligand binding to either the mGluR4 or the mGluR7 subunit induces some closure of unliganded mGluR2, with greater occupancy of the mGluR2 closed conformation in the presence of the mGluR7 subunit. (D) Representative traces for conditions shown in panel (C).

We next asked whether cooperative interactions between subunits could directly drive clamshell closure, even in the absence of a ligand capable of stimulating closure of a given subunit. We were motivated by our previous observation that mGluR2/7 fully occupies an active conformation with agonist binding to only the mGluR2 subunit or to the mGluR7 subunit (*13*). We tested this possibility by monitoring mGluR2 clamshell closure in mGluR2/7 and mGluR2/4, in the presence of the group III–selective agonist LSP4-2022 (*32*, *33*) to ensure that only mGluR7 or mGluR4, but not mGluR2, were bound by ligand. Indeed, LSP4-2022 induced closure of the mGluR2 clamshell in mGluR2/4 and, to a larger extent, in mGluR2/7, as seen by some occupancy of the high-FRET state of the mGluR2 clamshell (Fig. 5C, Extended Data Fig. 6). The larger influence of mGluR7 over mGluR4 is consistent with the greater positive cooperativity in mGluR2/7 (*13*, *23*). These observations show that activation of a single subunit can influence ligand binding to, and activation of, the empty partner subunit.

## Discussion

Homomeric and heteromeric combinations of the eight mGluRs differentially detect and respond to glutamate across a range of temporal and spatial scales (*34*, *35*). Here, we investigate homodimeric and heterodimeric versions of a prototypical presynaptic mGluR, mGluR2, but the multistate nature of mGluR activation appears to extend across the mGluR family and to other Family C GPCRs as well (*36*). Our experimental and computational analyses reveal that the diversity of responses to ligand across mGluRs emerges, at least in part, from differences in the dynamics and cooperativity of LBD clamshell closure. This early event in the activation pathway is coupled to LBD lower-lobe twisting, but the coupling is loose, such that full clamshell closure yields only partial occupancy of the twisted orientation of the LBD lower lobes. Likewise, coupling between the twisting of the LBD lower lobes and approach of the two CRD linkers is also loose, such that conditions that give rise to complete twisting of the LBD lower lobes (e.g., addition of a positive allosteric modulator or of heterotrimeric Gi) result in mixed populations of low, medium-, medium-high and high-FRET states of the CRD linkers. Strikingly, the high-FRET conformation, whose CRD–CRD separation distance approximately matches distances seen in Gi-bound cryo-EM structures, represents a very small fraction of the glutamate bound state, suggesting infrequent transitions to the conformation that can bind Gi, likely slowing the initiation of signaling and leaving considerable headroom for positive allosteric modulation of the kind seen with BINA. Strikingly, even though the CRDs are so dynamic, binding of BINA or Gi stabilizes the LBD-twisted state, which serves to maintain a degree of LBD-CRD coupling. Strikingly, even though the CRDs remain dynamic, coupling of transmembrane and intracellular regions to the LBD is maintained, as BINA and Gi stabilize the LBD-twisted conformation.

Our smFRET measurements and MD simulations reveal the existence of an mGluR2 intermediate in which the lower lobes of the LBD are partially relaxed away from each other despite the fact that both clamshells are closed around ligand. This inactive intermediate readily interconverts with an active LBD conformation, and our simulations reveal interactions at the dimerization interface that stabilize the active conformation. Recent cryo-EM studies of a post-synaptic, Gq-coupled mGluR, mGluR5, revealed a similar, agonist-bound intermediate conformation (Extended Data Fig. 7) (*15*) in which the lower lobes have partially relaxed towards their inactive orientation, consistent with a recent proposed model for activation of mGluR (*20*). We propose that sequence variation across the dimer interface in mGluR homo- and heterodimers can alter the relative stability of this intermediate to affect signaling efficacy. In MD simulation, the interactions that favor twisting and association of the LBD lower lobes are well conserved across pre-synaptic (Group II and Group III) mGluRs but may be specifically modulated in mGluR7-containing heterodimers (*5*), where an arginine substitution of neighboring serine S176 (mGluR2 numbering) in the R177-containing dimerization interface helix may alter packing interactions with that helix, and/or in Group I/II heterodimers, where Group I mGluRs possess hydrophobic residues in place of R177. Notably, a coevolution-based analysis of mGluR sequences identified the R177 position as a site of high co-variation with residues in the binding pocket, in agreement with our observation of coupling between the binding pocket and the R177 helix (*37*). We propose that stabilization of the R177-containing helix D by glutamate, as seen in HDX-MS, ensures proper alignment of the lower LBD dimer interface to form the contacts that stabilize the activated conformation and gives rise to cooperativity between the subunits.

Our smFRET data provide evidence that mGluRs can populate an ensemble of conformations whose occupancy is modulated by orthosteric agonist. For example, subtype-specific agonist binding to the clamshell of one subunit can drive closure of the empty partner mGluR subunit in Group II/III heterodimers, resulting in a variety of clamshell conformations (O/O, C/C, and O/C). Meanwhile, the LBDs can twist between relaxed and active conformations (R or A), and these LBD states are non-rigidly coupled to at least four conformations of the CRD. Low occupancy of the closest proximity (high-FRET) state of the CRD, which is favored by Gi binding, is likely rate-limiting for signaling onset and dampens efficacy, thereby providing considerable head-space for positive modulation by lipid environment, nanobodies and naturally occurring transsynaptic binding partners modulate mGluR function (*15*, *38–41*). Additional features outside of the LBD core, including within the disulfide-rich loop and the CRDs, further tune cooperativity in heterodimers (*24*, *42*, *43*), giving rise to a broad diversity of conformational responses to ligand. Heterodimers appear capable of modulating the coupling between ligand binding pocket closure and adoption of the Gi-coupled, active state by directly altering the fraction of the time the clamshell of either subunit populates a closed conformation. These observations suggest a framework for the design of allosteric modulators that can act within the extracellular dimerization interface to specifically target and tune the properties of mGluR heterodimers.

## Acknowledgements

We thank Stephen Brohawn, Kaavya Krishna Kumar, Antoine Koehl, Kevin Larsen, Shimon Yudovich, Joy Zhong and all members of the Isacoff and Marqusee labs for assistance and helpful discussion. We thank Betsy White for purification of heterotrimeric Gi1. Anton 2 computer time was provided by the Pittsburgh Supercomputing Center (PSC) through grant R01GM116961 from the National Institutes of Health. The Anton 2 machine at PSC was generously made available by D.E. Shaw Research. Support was provided by the National Institutes of Health (R01NS119826 and 1RF1MH123246 to E.Y.I.; K99GM148823 to N.L.; R35GM149319 to S.M.), a UC Berkeley Miller Postdoctoral Fellowship (N.L.) and a 2023 McKnight Pecot Fellowship (S.S.). S.M. is a Chan Zuckerberg Biohub Investigator. E.Y.I. is a Weill Neurohub Investigator.

## Author Contributions

N.L., C.H., S.M. and E.Y.I. conceived of the project. N.L., S.M. and E.Y.I. wrote the manuscript, with feedback from all of the authors. N.L. performed the experiments, with help from C.H. and J.B. on smFRET and guidance from S.M. on HDX-MS. N.L. and S.S. performed and analyzed the MD simulations. N.L. performed the analysis. C.S. helped with the molecular biology.

## Data and materials availability

smFRET data, HDX-MS data, simulation trajectories, and analysis code will be deposited in Zenodo prior to publication of the manuscript.

## Methods

### Constructs

We developed several mGluR2 constructs for FRET analysis using the hemagglutinin (HA) epitope-SNAP-tagged rat mGluR cloned into the pRK5 backbone described previously (*13*). cDNAs for rat mGluR2, mGluR3, mGluR4 and mGluR7 were amplified to include an N-terminal GSGS linker and an mluI cut site. These sequences were then inserted into the base construct using mluI and xbaI restriction enzymes, resulting in constructs composed of the mGluR5 signal peptide fused to the HA epitope, followed by a GSGS linker, and then by the mGluR cDNA. To monitor LBD closure, we removed the HA epitope and GSGS linker and introduced rare codon TAG at sites corresponding to Gln359 and Ser463. To monitor intersubunit twisting, we retained the HA epitope and GSGS linker and introduced the rare codon TAG into site Ala248 (for the lower lobe) and Ala548 (for the CRD linker). Various point mutations were introduced using site-directed quick-change mutagenesis. The construct encoding the tRNA/tRNA synthetase pair, Mm-PylRS-AF/Pyl-tRNACUA, was a gift from Howard Hang (Addgene plasmid #122650; http://n2t.net/addgene:122650; RRID:Addgene_122650). We introduced the following modifications: insertion of a nuclear export signal (*44*), a post-transcriptional regulatory element (*45*), three additional copies of the tRNA, and a translation elongation factor.

For protein purification of isolated mGluR2 LBD, we ordered codon-optimized mGluR2 (residues 22–502) from Twist Biosciences, which we inserted into the backbone of pcDNA-Zeo-tetO containing a hemagglutinin signal peptide fused to a FLAG (DYKDDDDK) epitope, kindly provided by Brian K. Kobilka (*15*).

### Cell culture and transfection

HEK-293T cells were obtained from the UC-Berkeley Cell Culture Facility and maintained in DMEM with 10% FBS in T-25 flasks. Prior to transfection, cells were seeded on poly-L-lysine coated 6-well plates at 10% confluence. Cells were transfected between 24 and 48 hours before harvesting. 100 mM trans-cyclooct-2-en-L-lysine (TCO*A) was diluted in 1 M HEPES to 20 mM and added to media for a final concentration of 250 µM immediately prior to transfection. To express the mGluR2 heterodimers for monitoring LBD closure, cells were transfected with 0.2 µg HA-GSGS-tagged mGluR2, 2.5 µg mGluR2-359TAG-463TAG, and 2.5 µg of Amber suppressor DNA, per well in a 6-well plate, using lipofectamine 3000. To express mGluR2 homodimers for monitoring intersubunit twisting, cells were transfected with 0.5 µg of HA-GSGS-tagged mGluR and 0.5 µg of Amber suppressor DNA using lipofectamine 2000.

For mGluR2 LBD expression, Expi293 GNTI– cells were obtained from the UC-Berkeley Cell Culture Facility and maintained in Expi293 expression medium in 125–250 mL flasks. Prior to transfection, cells were split to a density of 1×10^6^ cells / mL and grown for 24 hours to > 2×10^6^ cells. Cells were transfected with DNA at a concentration of 1 µg / mL of culture volume using a 4:1 ratio of PEI to DNA combined in hybridoma serum-free media. Valproic acid was added at a final concentration of 2.2 mM 16-24 hours after transfection. Cultures were grown for up to five days after transfection before collection of media for protein purification.

### Purification of mGluR2 LBD for hydrogen–deuterium exchange

After five days of protein expression, media was collected by spinning cells at 2000 rpm for 20 min at 4 C. Media was then filtered using a 0.2 µm filter, and a gravity column of 1 mL of resuspended anti-FLAG m2 resin was equilibrated with 50 mL of buffer (100 mM NaCl, 20 mM HEPES, pH 7.5). 250 mL of media were applied to the gravity column. The column was washed with 40 mL of buffer. Sample was eluted using 3xFLAG peptide diluted in wash buffer to a concentration of 150 µg / mL in 1 mL increments. Protein was concentrated and snap frozen prior to size exclusion chromatography. Protein was diluted to 500 ul in buffer and run over a Superdex 200 10/300 increase column (Cytiva). Protein eluted as a single peak at its expected dimer molecular weight of 107 kDa. Protein was concentrated to 9.7 µM (monomer) and snap frozen.

### smFRET sample preparation

On the day of each imaging experiment, cells were washed with extracellular (Ex) solution containing 10 mM HEPES, 135 mM NaCl, 5.4 mM KCl, 2 mM CaCl_2_, 1 mM MgCl_2_, pH 7.4. Pyrimidyl-tetrazine-AF555 and pyrimidyl-tetrazine-AF647 (Jena Biosciences) were diluted in EX buffer to 300 nM; biotinylated anti-HA polyclonal antibody stored at 0.5 mg / mL was added to the dye-containing EX solution to a final concentration of 0.5 µg / mL. Cells were incubated at 37 C for 15–20 minutes, then washed twice with Ex buffer. Cells were harvested and kept on ice and then spun down at 5,000 x g in a benchtop centrifuge cooled to 4 C for 5 minutes. Subsequently, cell pellets were lysed in lysis buffer (150 mM NaCl, 1 mM EDTA, with protease inhibitor) containing 1% MNG / 1% GDN / 0.1% CHS. After incubation for an hour at 4 C, lysate was centrifuged at 21,000 x g for 25 minutes. Supernatants were transferred to polycarbonate ultracentrifuge tubes for ultracentrifugation at 4 C for 45 min at 75,000 rpm to remove aggregates. Subsequently, supernatants were kept on ice prior to imaging. Note that all buffers were made with ultrapure reagents to eliminate trace glutamate.

### smFRET measurements

Imaging chambers with six to eight flow cells were constructed using passivated glass coverslips coated in mPEG / biotin-PEG, as previously described. Flow cells were washed with T50 (150 mM NaCl, 50 mM Tris-HCl, pH 7.5) buffer. Neutravidin antibody was diluted to 10 nM in T50 solution, applied to flow cells, and incubated for approximately 20 minutes. Flow cells were washed twice more with T50 buffer to remove excess neutravidin. Samples were then diluted 2x to 50x in Ex buffer containing 0.01% MNG / 0.01% GDN/ 0.001% CHS before being applied to coverslips and allowed to adhere for several minutes until reaching optimal surface density (∼800 molecules per imaging area). Flow cells were then washed extensively (up to 100x flow cell volume). Receptors were imaged in Ex buffer containing 5 mM trolox, 0.01% MNG / 0.01% GDN/ 0.001% CHS, and 2:1 protocatechuic acid/protocatechuate-3,4-dioxygenase (Millipore Sigma).

Samples were imaged with a 1.65 na, 60x objective (Olympus) on a TIRF microscope with 100 ms time resolution. We employed a 532 nm laser (Cobolt) for donor excitation, a 632 nm laser (Melles Griot) for acceptor excitation, and a Photometrics Prime 95B cMOS camera with a 100 ms acquisition time. Movies were typically 90 seconds in length, for a total of 9000 frames.

In some cases, ligands and/or G protein (heterotrimeric Gi) were added to imaging chambers. We obtained LY379268 from Tocris Biosciences and added it to imaging buffer at a final concentration of 20 µM; we also obtained biphenyl-indanone A (BINA) from Tocris Biosciences and added it to imaging buffer at a final concentration of 100 µM. Heterotrimeric Gi in DDM/CHS and ScFV antibody were gifts from the Kobilka laboratory and purified as described (*49*). Briefly, Gi was incubated in 0.02% DDM/CHS on ice for 30 minutes before addition of 0.5 µl apyrase (NEB) and ScFV at a 1:1 ratio (the final concentrations of Gi and ScFV were both ∼90 µM) to favor nucleotide-free G protein. Complex was incubated for one additional hour on ice, diluted 1:9 in imaging buffer with 0.02% DDM/CHS and added to chambers with Gi at final concentration of 4–5 µM and glutamate at 10 mM, incubated for 20 minutes, and then imaged.

### smFRET data analysis

We employed SPARTAN to analyze resulting smFRET movies (*46*). For the *gettraces* function, we used default values, but we set the Cy3-to-Cy5 crosstalk to 0.150, the donor/acceptor channel scaling to 1.0, autopicking sensitivity to 5.0 (occasionally, to 4.0), and the integration window size to 10.0. For the *autotrace* function, we again used default values, but we set the minimum FRET lifetime to 5.0 seconds. Finally, using the *sorttraces* function, we manually retained traces for further analysis that demonstrated anti-correlation in donor and acceptor fluorescence, exhibited relatively constant total fluorescence, and underwent single acceptor and donor bleaching events. Histograms were compiled for the first 5.0 seconds across all selected traces from each movie and exported from SPARTAN with a bin size of 0.02 for further analysis in Python. smFRET histograms shown in figures represent the average of at least three (but typically, 5) movies collected on the same day; error bars represent the standard error of the mean across those movies. For experiments involving addition of ligand, we typically analyzed data from movies collected 5–15 minutes after ligand addition. For experiments involving addition of Gi, we initiated imaging after incubation for 20 minutes on the chip. Experiments were repeated at least twice on separate days. To calculate the fraction of receptors in high-FRET states for dose-response curves, we first used ebFRET (*47*) in SPARTAN to determine histogram centers and standard deviations with three-state models (one state corresponds to zero FRET). We then fit histograms to sums of Gaussians using the scipy fit function in Python; for the lower-lobe twisting sensor, we used the ebFRET-determined histogram centers as fixed parameters, while for the CRD twisting sensor, we used the ebFRET-determined histogram centers only as initial guesses, as the low FRET peak shifts slightly with increasing glutamate concentration. Dwell times within each state were exported and plotted from ebFRET (Extended Data Fig. 5C).

### Molecular dynamics simulation

We initiated simulations of the mGluR2 ligand binding domain (LBD) from the crystal structure of human mGluR2 bound to the agonist (1S,2S,5R,6S)-2-aminobicyclo[3.1.0]hexane-2,6-dicarboxylic acid, also known as LY354740 or eglumegad (*29*). We retained co-crystallized chloride ions and waters. Prime (Schrödinger) was used to model hydrogen atoms and missing side chains; neutral acetyl and methylamide groups were used to cap protein termini. Note that the disulfide-containing loop extending from residues Ser111 to Pro133 were left unmodeled. We retained titratable residues in their dominant protonation state at pH 7.0, resulting in protonation of His49, with the exception of Asp188, which we retained in its charged state because of its instability in simulation when neutralized.

We used *tLeap* in AmberTools (2020) to prepare the mGluR2 LBD for simulation (*48*). We parameterized the co-crystallized agonist using *antechamber* with the General Amber Force Field 2 and ensured that the ligand retained a net charge of –1.0 in all simulations (*49*). We employed the four-point OPC water model, and the ff19SB protein force field (*50*, *51*). Water-box dimensions were chosen to maintain an 18 Å buffer between the protein image and the edge of the box, resulting in a box size of 125 Å × 125 Å × 125 Å. Sodium and chloride ions were added to neutralize the system to a concentration of 150 mM. Boxes were composed of 246,886 atoms (this count includes the ‘dummy’ atom employed in the four-point OPC water model).

We initiated simulations using the Compute Unified Device Architecture (CUDA) version of Particle-Mesh Ewald Molecular Dynamics (PMEMD) in AMBER on single graphical processing units (GPUs) (*52*). Simulations were performed using the AMBER18 software. Systems were first minimized using three rounds of steepest descent minimization followed by a conjugate gradient minimization step. Systems were heated from 0 to 100 K in the NVT ensemble over 12.5 ps and then heated from 100 K to 310 K in the NPT ensemble over 125 ps at 1 bar, with 10.0 kcal mol^-^_1_Å^2^ harmonic restraints placed on non-hydrogen protein atoms, ligand atoms, and co-crystallized ions. Systems were then equilibrated at 310 K in the NPT ensemble at 1 bar in 2-ns increments, with harmonic restraints tapered by 1.0 kcal mol^-1^Å^2^ for 10 ns and then by 0.1 kcal mol^-1^Å^2^ for 20 additional nanoseconds, for a total of 30 ns of equilibration. Production simulations were carried out in the NPT ensemble at 310 K and 1 bar, using a Langevin thermostat for temperature coupling and a Berendsen barostat with isotropic control for pressure coupling. We applied hydrogen mass repartitioning in order to employ a 4-fs time step, and we constrained bond lengths to hydrogen atoms using SHAKE (*53*). Non-bonded interactions were cut off at 9.0 Å; long-range electrostatic interactions were calculated using Particle Mesh Ewald (PME) with an Ewald coefficient of 0.30768 and a B-spline interpolation order of 4. The FFT grid size was chosen such that the width of each grid cell was ∼1 Å. Trajectory snapshots were saved every 200 ps. Production simulations on Savio were visually checked for stability prior to transfer of the simulation to an Anton 2 machine.

To initiate simulations on Anton 2, we transferred the system parameter file (.prmtop) and an ASCII-readable restart file containing velocities from the end of the first production step calculated on Savio (typically, 60 ns of production after removal of harmonic restraints). Simulations were approximately 15.0 µs in length and employed a RESPA integrator, with time steps of 4.0 fs and long-range interactions calculated every two steps. These simulations employed an MTK barostat, a Nose-Hoover barostat, and isotropic pressure control. Simulation snapshots were saved every 240 ps. Simulations were downsampled further to 12 ns and stripped of waters to reduce file size for analysis, unless indicated otherwise.

Simulation analyses were performed using Visual Molecular Dynamics (VMD) and visualized using the PyPlot package from Matplotlib. To measure LBD opening, we calculated the distance between the Cα atoms of residues Tyr144 and Ser272. To measure LBD separation, we calculated the distance between the centers of mass of the two lower lobes, each composed of residues 188– 317 and 452–474 of each protomer. To assess hydrogen bonding interactions between residues at the dimerization interface, we used VMD’s *hbonds* function, with a donor–acceptor atom cutoff of 3.5 Å and a donor–hydrogen–acceptor angle cutoff of 50°. For sets of interactions involving atoms from the same pairs of residues, a residue–residue interaction was considered present if any one of those pairs was interacting. To calculate root-mean-square fluctuations for helix D, adjacent to Arg177, we aligned simulation frames on residues 155–166 and 180–188, which flank the region of interest, for either chain. We averaged simulation frames from two Anton simulations that did not transition to a relaxed-closed/closed (R-C/C) intermediate to generate the average structure and then calculated the root-mean-square fluctuation for the Cα atoms of residues 166–180 using 200 pre-transition frames or 200 post-transition frames for each simulation that transitioned to an R-C/C intermediate.

### Hydrogen–deuterium exchange monitored by mass spectrometry

Purified mGluR2 LBD was diluted slightly (9:1) to 8.7 µM (monomer) in buffer (100 mM NaCl, 20 mM HEPES with or without monosodium glutamate at a 10x concentration of 100 mM, pH 7.5, and allowed to incubate with ligand for 20 minutes. Deuterated buffer was prepared by resuspending NaCl to 100 mM and HEPES to 20 mM (and, for the glutamate-bound condition, monosodium glutamate to 10 mM) with D_2_O (Sigma Aldrich) and adjusted to pH_read_ = 7.3 using NaOD (Sigma Aldrich). To initiate exchange, samples were diluted 1:10 into D_2_O buffer and quenched on ice with a 2X quench buffer (3 M urea, 20 mM TCEP, pH 2.4) for one minute prior to flash freezing in liquid N_2_. Samples were stored at –80 C prior to LC/MS analysis.

Samples were thawed and injected into a valve system cooled to 2 C (Trajan LEAP) coupled to a Thermo Ultimate 3000 LC, with buffer A (0.1% formic acid) flowing at 200 µl/min. Peptides were subjected to proteolysis via two in-line protease columns manually packed with porcine pepsin or aspergillopepsin (held at 10 C) and desalted onto a trap column (1 mM inner diameter x 2 cm, IDEx C-128) manually packed with POROS R2 reversed-phase resin (Thermo Scientific). Peptides were then separated onto a C18 analytical column (Waters Acquity UPLC BEH, pore size 130 Å, particle size 1.7 µm, 2.1 mm ID X 50 mm) with buffer flowing at a rate of 45 µl/min and buffer B increasing in concentration from 5% to 40% over the first 14 minutes and from 40% to 90% over the next 30 s. Two sawtooth gradients to wash the analytical column were performed before the column was equilibrated back to 5% buffer B prior to the next injection. Peptides were eluted into a Q Exactive Orbitrap Mass Spectrometer (Thermo Fisher) operating in positive ion mode (MS1 settings: resolution 140000; AGC target 3e6; maximum IT 200 ms; scan range 300-1500 m/z). Tandem mass spectrometry analysis were carried out with MS1 settings same as above but with resolution of 70000 and MS2 settings of: resolution 17500; AGC target 2e5; maximum IT 100 ms; loop count 10; isolation window 2.0 m/z; NCE 28; and charge states of 0, 1 and >8 excluded, with dynamic exclusion of 15.0 s.

Over 700 peptides, including 127 glycosylated peptides, were identified from MS2 data using Byonic (Protein Metrics). Deuterium uptake was analyzed using HD-Examiner (Version 3.1, Sierra Analytics) using default settings after adjusting for 90% maximal deuteration of all exchanged samples. Deuterium uptake information was exported from HDExaminer for further analysis with Python.

---

**Extended Data Fig. 1.**
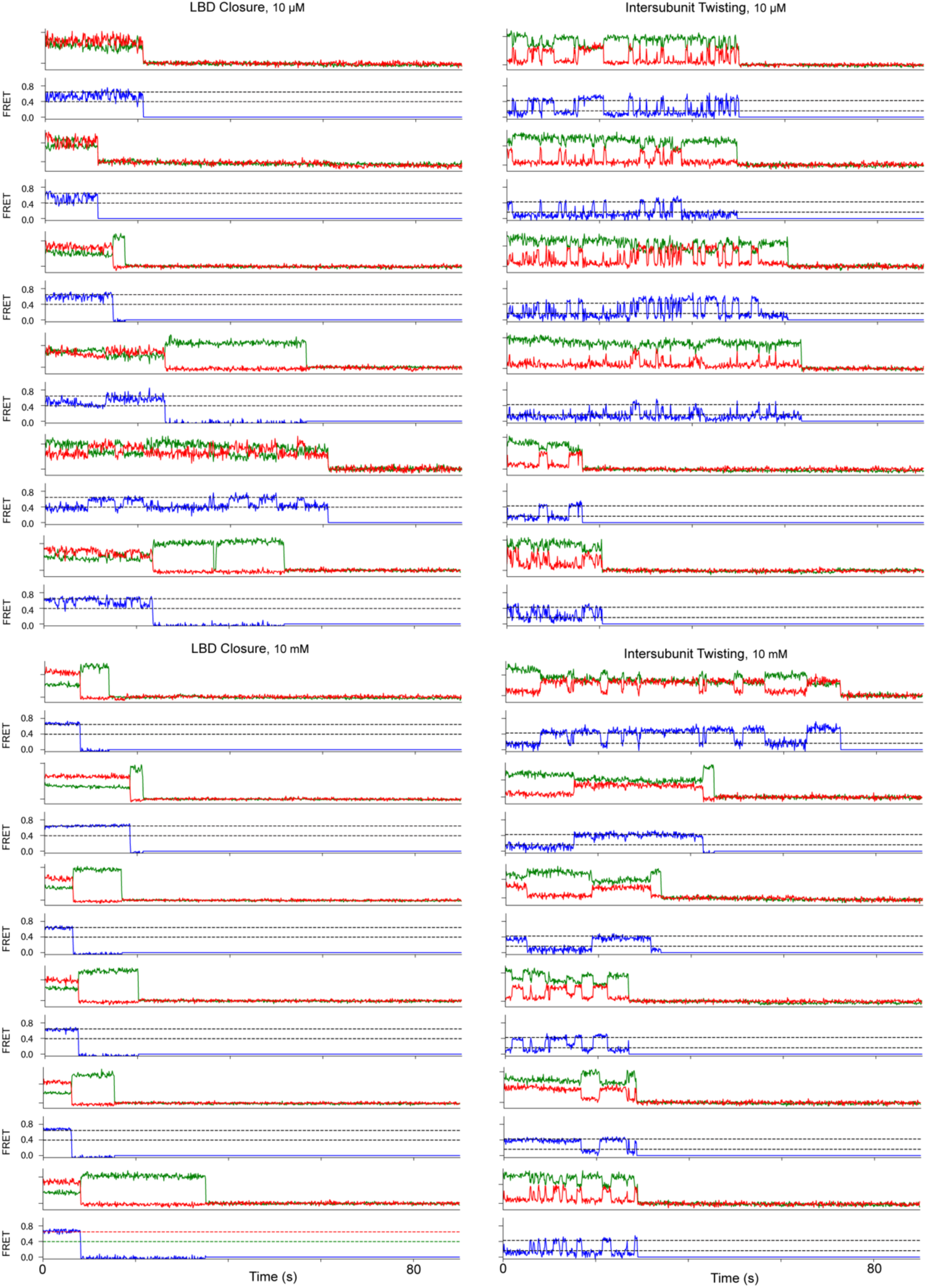
Representative smFRET traces. Donor (green) and acceptor (red) intensities (top) and FRET values (blue; bottom) for the LBD closure (left) and inter-subunit twisting (right) FRET pairs.

**Extended Data Fig. 2.**
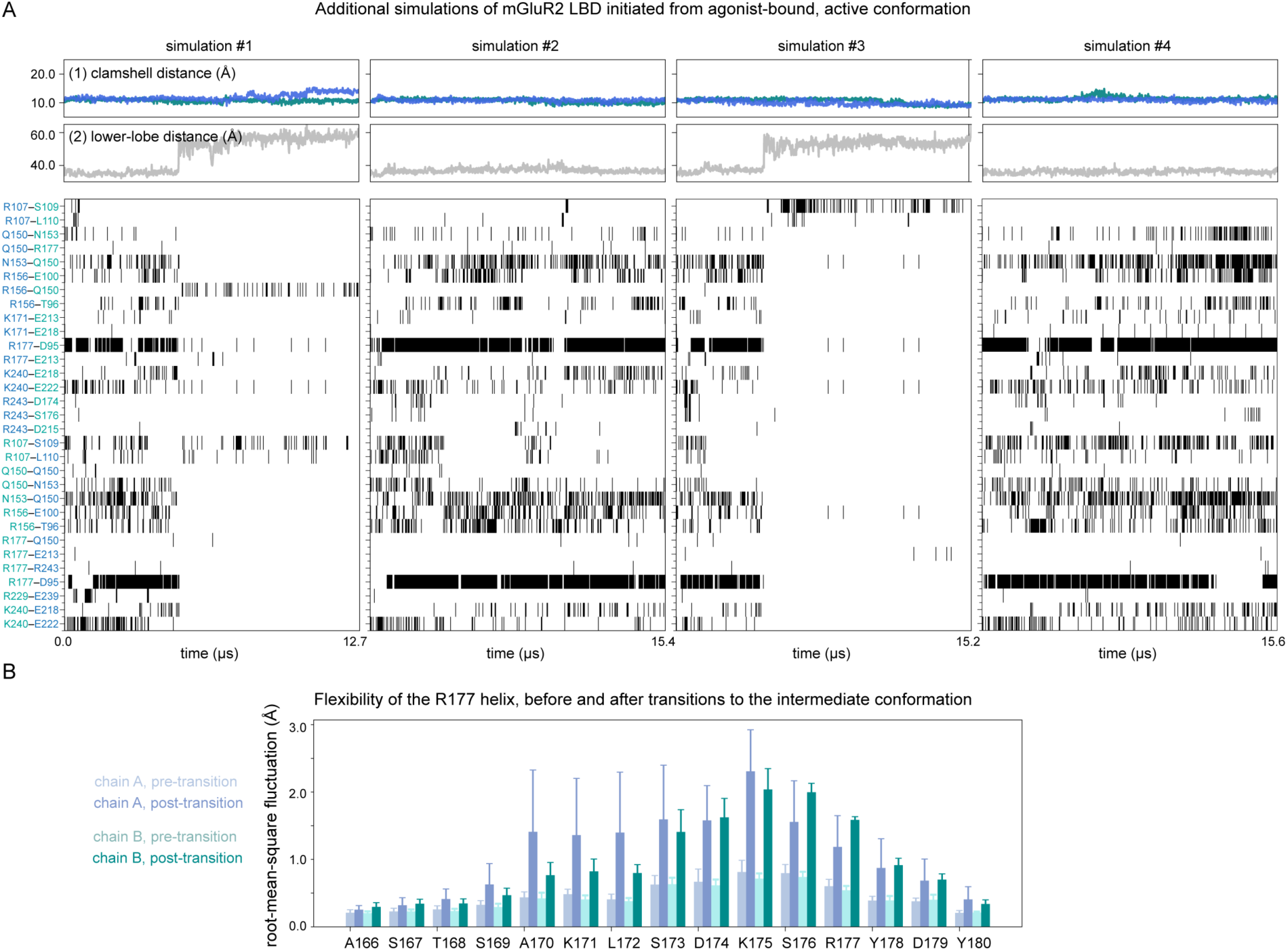
(A) Traces and interface interaction time courses for additional simulations. Simulations 1 and 3 show transitions to a relaxed, LBD-closed intermediate, and data for simulation 5 is represented in the main text figure. (B) Changes in the flexibility of the R177-containing helix, before and after the transition (*n* = 3 simulations; error bars represent s.e.m.). RMSF analysis carried out on 2 µs pre- and post-transition; average structure calculated from simulations 2 and 4, which do not transition away from initial state (left).

**Extended Data Fig. 3.**
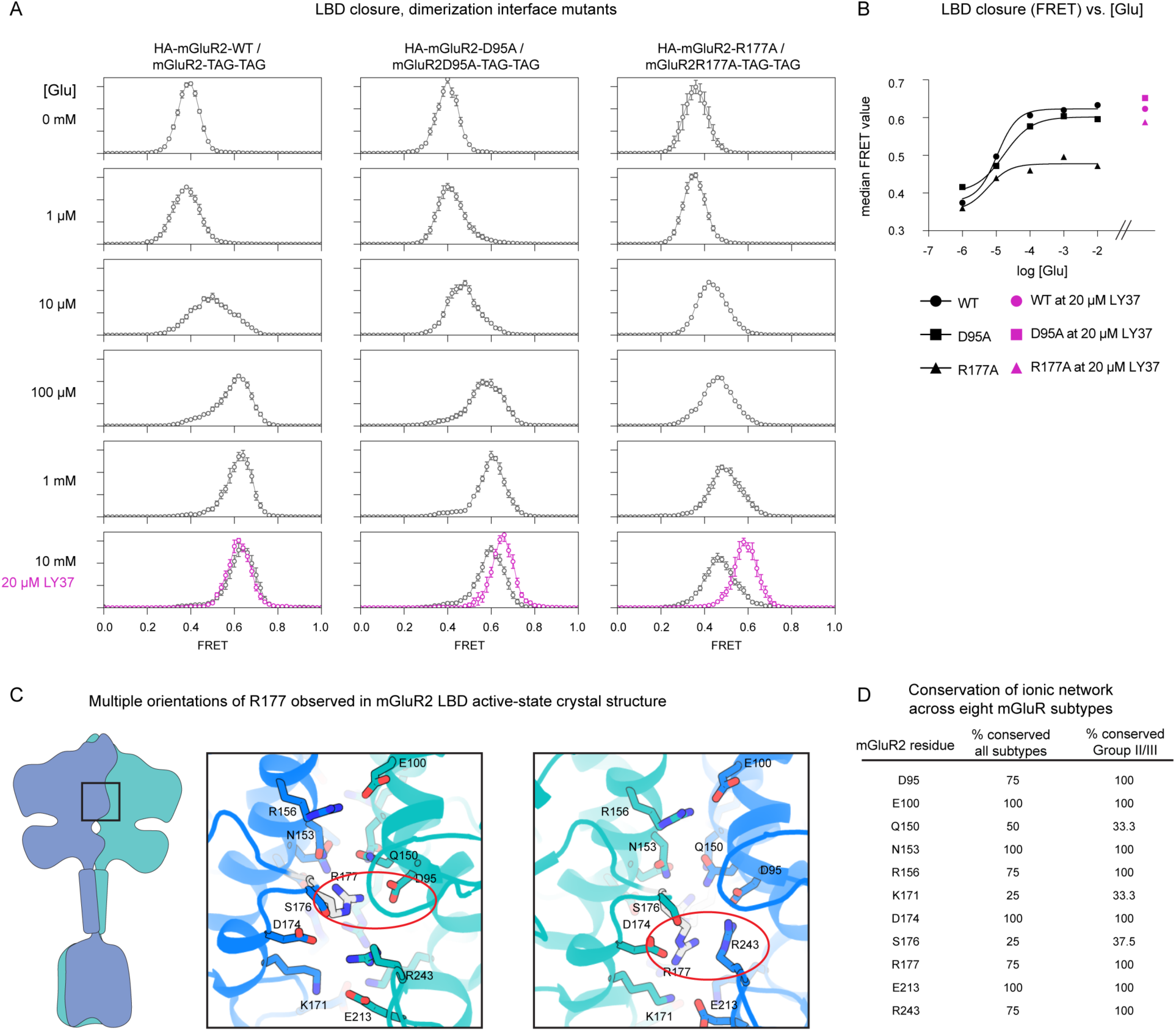
(A) smFRET distributions for LBD closure for two mutants demonstrate that R177A, but not D95A, leads to a reduction in glutamate affinity, but that in the presence of a strong agonist, both mutants populate fully closed states. (B) These changes are summarized by the plot on the right, which shows the median FRET value for each averaged smFRET distributed plotted vs. glutamate concentration. Pink circles represent data collected in presence of 20 µM LY37. (C) In a crystal structure (PDB entry 4XAQ) of the mGluR2 LBD, R177 adopts two distinct orientations. We speculate that the non-equivalent effects of D95A and R177A on ligand affinity and intersubunit twisting are due, at least in part, to the ability of R177 to engage R243 on the opposite protomer via a π–π stack, one of the two R177 orientations observed in the crystal structure of agonist-bound mGluR2 LBD (29). Classical molecular mechanics forcefields do not explicitly represent π–π interactions, suggesting that our simulations may over-represent the orientation of R177 that engages D95. Additionally, introduction of R243A in single-molecule constructs substantially reduced expression, preventing smFRET investigation. (D) Conservation of the interface network across the eight mGluR subtypes found in *R. norvegicus*.

**Extended Data Fig. 4.**
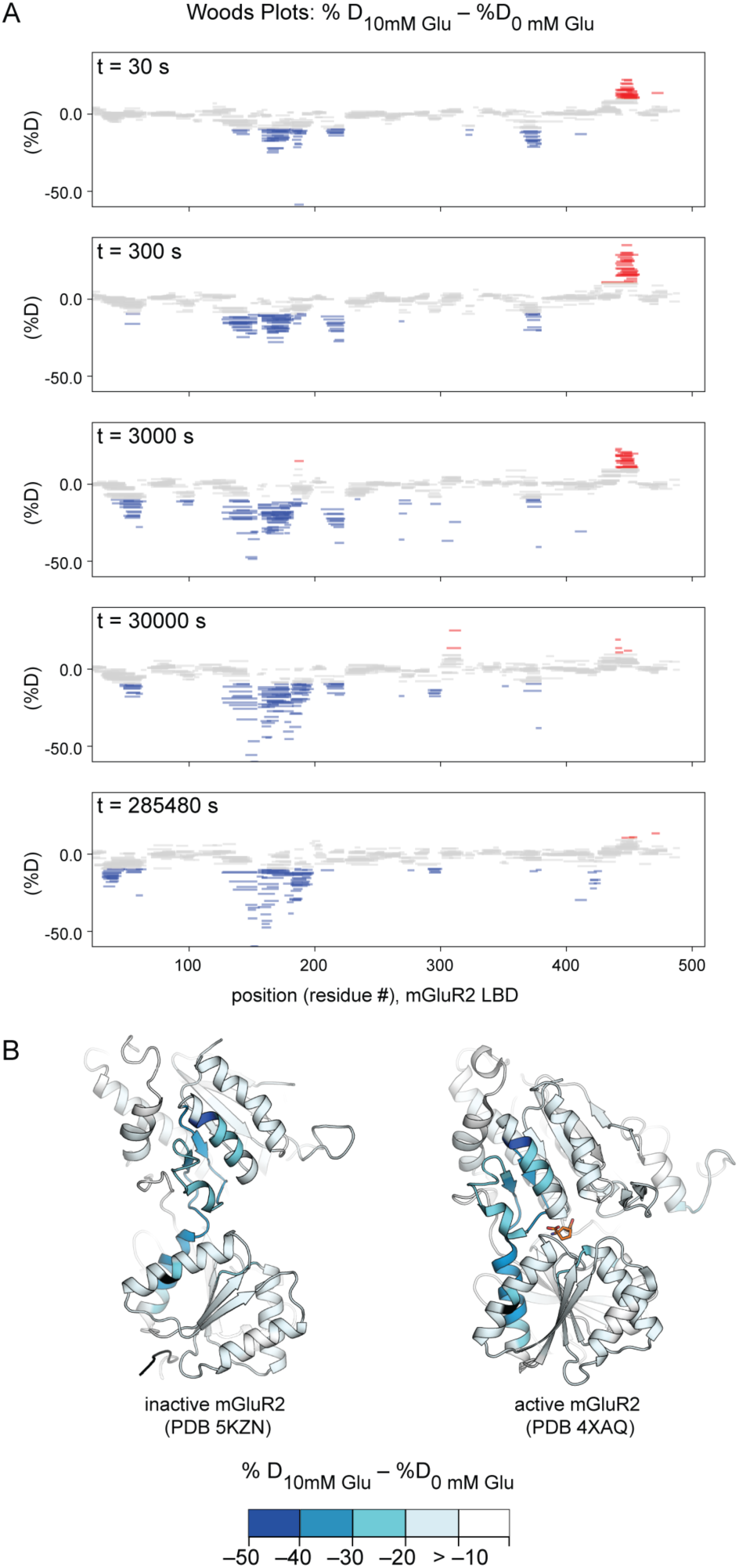
(A) Woods plots showing (top) change in hydrogen exchange protection vs. peptide sequence position at increasing exchange times. Peptides are colored blue if they exhibit an *increase* in protection of at least 10% in the presence of glutamate or red if they exhibit a *decrease* in protection of at least 10% in the presence of glutamate. (B) Dimerization interfaces of two mGluR2 crystal structures are colored by the maximal change in protection for any peptide covering a given residue position.

**Extended Data Fig. 5.**
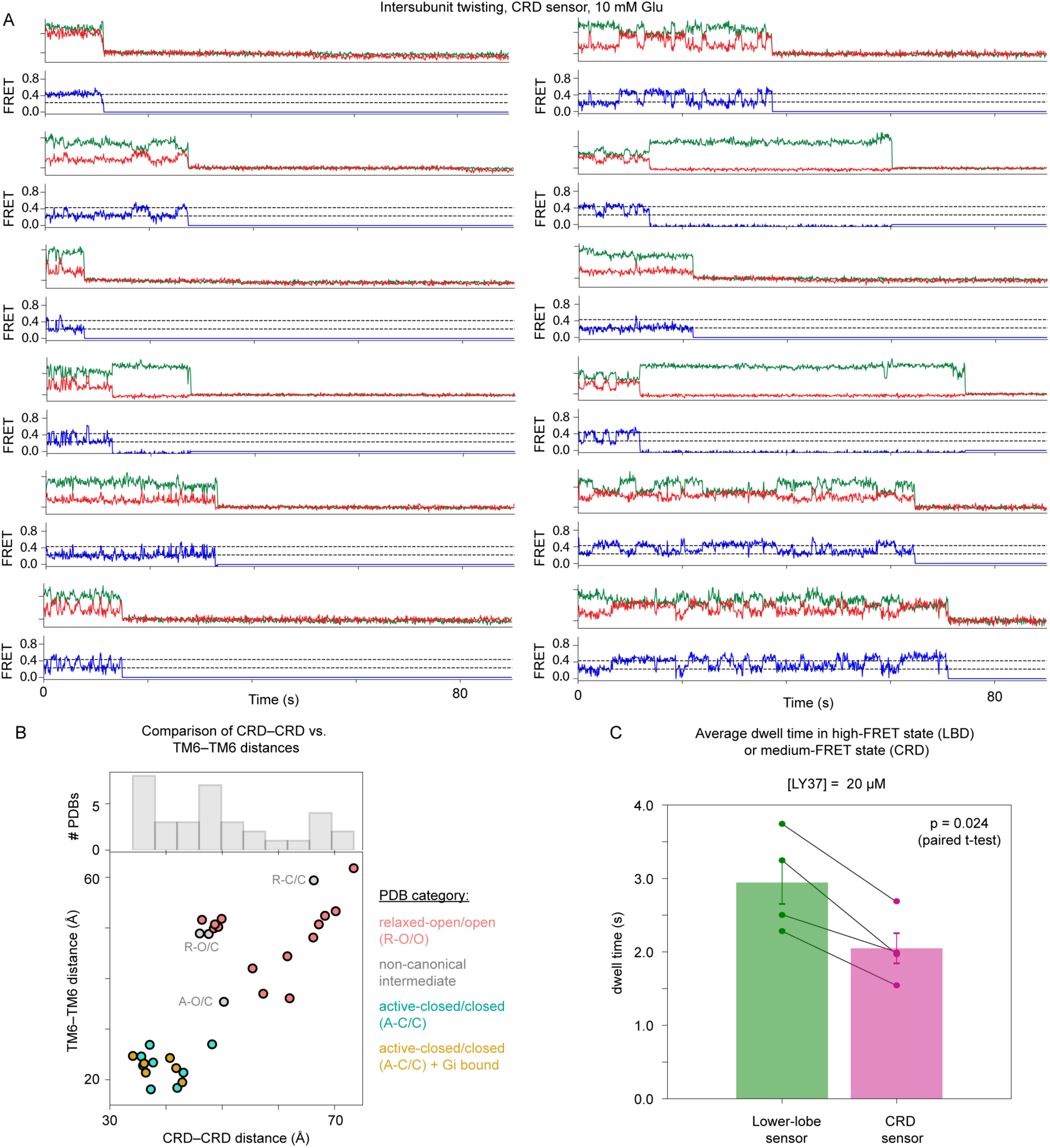
(A) Representative smFRET traces for the CRD reporter, collected in the presence of 10 mM glutamate (left). (B) Scatter plot of TM6–TM6 vs. CRD–CRD distances across different full-length cryo-EM structures of mGluRs. A histogram of CRD–CRD distances reveals the heterogeneous spectrum of CRD–CRD distances observed across mGluR homo- and heterodimers. Cα positions for Ala548 (mGluR2) or the equivalent position in other mGluRs were used to determine CRD–CRD distances; Cα positions for Phe756 (mGluR2) or equivalent in other mGluRs were used to determine TM6–TM6 distances (C) Dwell times calculated from a two-state HMM fit carried out using ebFRET in SPARTAN. Each point corresponds to the average dwell time from one of four separate experimental days; dwell times for each sensor were compared on each day.

**Extended Data Fig. 6.**
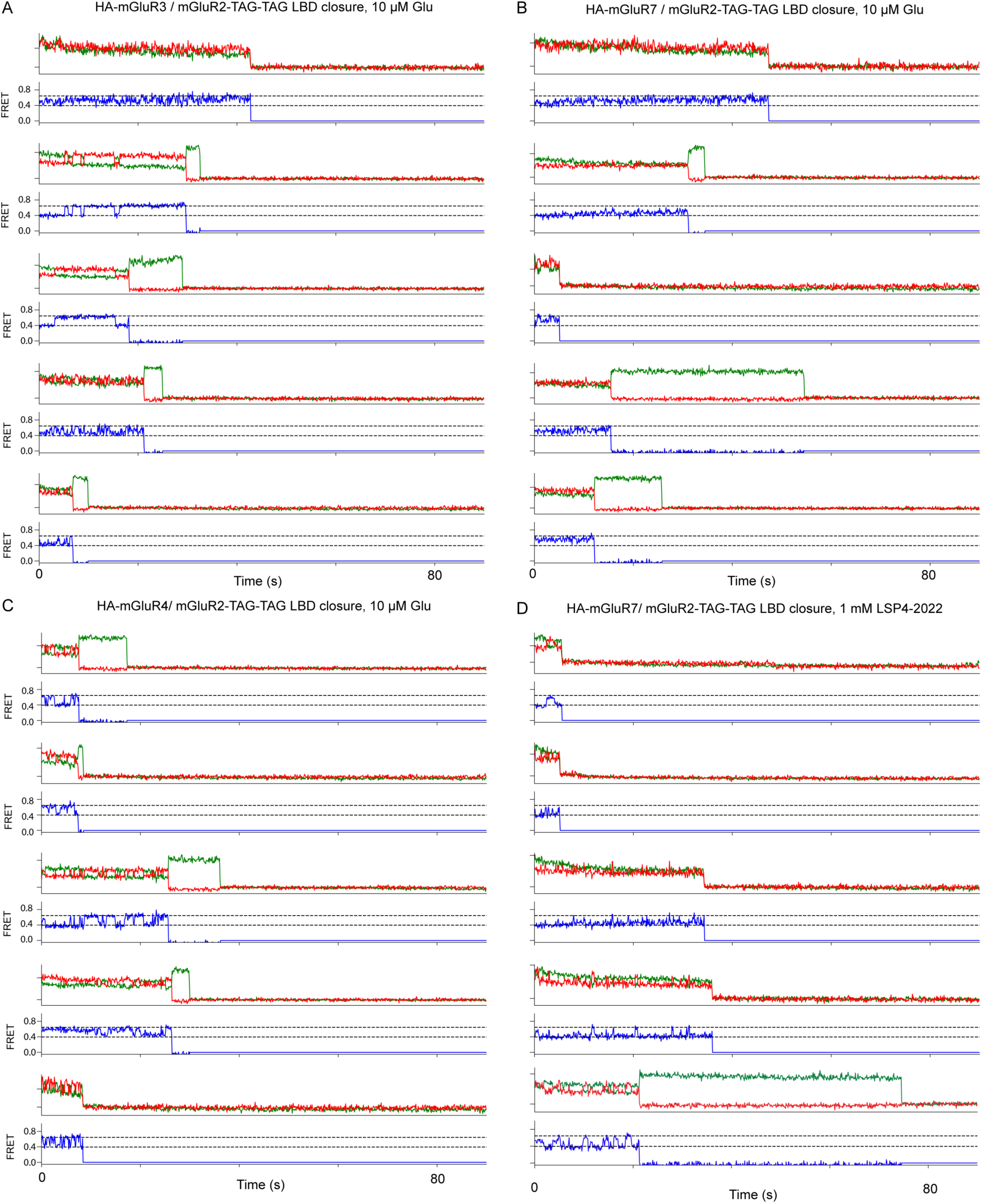
smFRET traces for the LBD closure reporter, which reveal differences in kinetics of transitions between the low- and high-FRET states. Specifically, the mGluR2 subunit adopts longer lived low- and high-FRET states in the mGluR2/3 and mGluR2/4 heterodimers (A, B) vs. the mGluR2/7 heterodimer (C), where those states are not easily resolved. In the presence of a Group III (mGluR7)- specific agonist (D), the mGluR2 subunit in mGluR2/7 heterodimers occasionally makes excursions to high-FRET states, indicating that mGluR2 is allosterically influenced by mGluR7.

**Extended Data Fig. 7.**
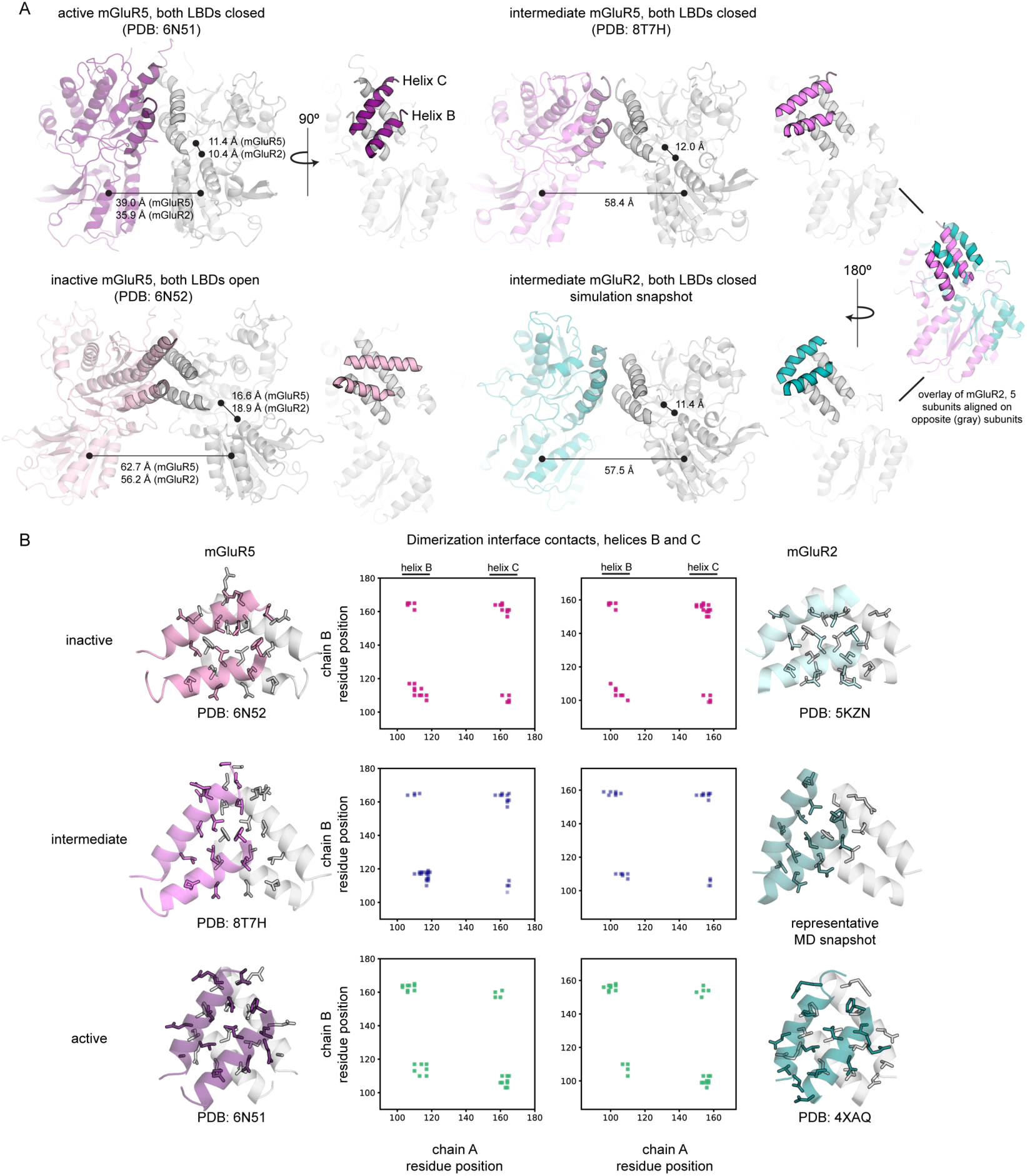
Comparison to structures of mGluR5 determined via cryo-EM. (A) The cryo-EM–captured mGluR5 intermediate (top, right) and the MD-determined mGluR2 intermediate (middle, right) both exhibit two closed LBDs with substantial separation between their lower lobes (≥ 50 Å). mGluR5 structures were compared to the mGluR2 active state structure (PDB 4XAQ) and the mGluR2 inactive state structure (PDB 5KZN). The distances between upper and lower lobes of each clamshell correspond to the Cα–Cα distance between residues 144 and 272 (mGluR2) or residues 151 and 280 (mGluR5). LBD distances correspond to the distance between either chain’s lower-lobe center of mass (residues 188–317 and 452–474 for mGluR2 or residues 195–327 and 465–487 for mGluR5). Note that simulated conformations of the mGluR2 LBD differ from that of the mGluR5 intermediate in terms of the relative positioning of the hydrophobic interface residues (overlay, far right) and in terms of (B) van der Waals contacts between upper lobe dimerization interface helices B and C. Contact matrices for six different mGluR5 (purple) and mGluR2 (teal) structures reveal conformation–specific contact patterns. In the R-O/O conformation, helices B and C form numerous self-contacts across the interface, while in the A-C/C conformation, helices B and C form numerous cross-interface interactions (off-diagonal elements). The simulated mGluR2 intermediate and the mGluR5 relaxed-closed/closed intermediate state are similar in that cross-protomer contact between helices B and C is minimal but differ in the degree to which one subunit has fully rotated with respect to the other, giving rise to the asymmetry observed in the mGluR2 intermediate’s contact matrix.

**Extended Data Table 1.** Counts of particles in smFRET experiments. 359-463-TAG corresponds to the clamshell sensor; 248-TAG corresponds to the lower-lobe LBD twisting sensor; and 548-TAG corresponds to the CRD sensor. * indicates conditions for which we did not manually sort traces.

| Figure | Sensor | Ligand/Condition | # Movies | Total particles |
| --- | --- | --- | --- | --- |
| Figure 1 | 248-TAG | 0 mM Glu | 5 | 163 |
| Figure 1 | 248-TAG | 1 uM Glu | 5 | 109 |
| Figure 1 | 248-TAG | 10 uM Glu | 5 | 106 |
| Figure 1 | 248-TAG | 100 uM Glu | 5 | 166 |
| Figure 1 | 248-TAG | 1 mM Glu | 5 | 181 |
| Figure 1 | 248-TAG | 10 mM Glu | 5 | 158 |
| Figure S6 | 248-TAG | 10 mM Glu + 100 uM BINA | 5 | 203 |
| Figure S6 | 248-TAG* | 10 mM Glu + 4 uM Gi | 5 | 560 |
| Figure 1 | 359-463-TAG + HA-GSGS-mGluR2 | 0 mM Glu | 7 | 242 |
| Figure 1 | 359-463-TAG + HA-GSGS-mGluR2 | 1 uM Glu | 6 | 162 |
| Figure 1 | 359-463-TAG + HA-GSGS-mGluR2 | 10 uM Glu | 6 | 144 |
| Figure 1 | 359-463-TAG + HA-GSGS-mGluR2 | 100 uM Glu | 6 | 172 |
| Figure 1 | 359-463-TAG + HA-GSGS-mGluR2 | 1 mM Glu | 6 | 155 |
| Figure 1 | 359-463-TAG + HA-GSGS-mGluR2 | 10 mM Glu | 6 | 155 |
| Figure 2 | 248-TAG | 20 uM LY37 | 4 | 186 |
| Figure 2 | 248-TAG, R177A | 20 uM LY37 | 4 | 344 |
| Figure 2 | 248-TAG, D95A | 20 uM LY37 | 4 | 536 |
| Figure 2 | 248-TAG, Q150A* | 20 uM LY37 | 4 | 479 |
| Figure 2 | 248-TAG, N153A* | 20 uM LY37 | 4 | 307 |
| Figure 2 | 248-TAG, R156A* | 20 uM LY37 | 4 | 453 |
| Figure 2 | 248-TAG, E218A* | 20 uM LY37 | 4 | 874 |
| Figure 2 | 248-TAG, E222A* | 20 uM LY37 | 4 | 713 |
| Figure 2 | 248-TAG, K240A* | 20 uM LY37 | 4 | 272 |
| Figure 3, S3 | 359-463-TAG R177A + HA-GSGS-mGluR2-R177A | 0 mM Glu | 5 | 266 |
| Figure 3, S3 | 359-463-TAG R177A + HA-GSGS-mGluR2-R177A | 10 mM Glu | 5 | 171 |
| Figure 3, S3 | 359-463-TAG R177A + HA-GSGS-mGluR2-R177A | 20 uM LY37 | 5 | 187 |
| Figure 4 | 548-TAG | 0 mM Glu | 4 | 339 |
| Figure 4 | 548-TAG | 1 uM Glu | 5 | 495 |
| Figure 4 | 548-TAG | 10 uM Glu | 5 | 585 |
| Figure 4 | 548-TAG | 100 uM Glu | 5 | 513 |
| Figure 4 | 548-TAG | 1 mM Glu | 4 | 392 |
| Figure 4 | 548-TAG | 10 mM Glu | 4 | 295 |
| Figure 4 | 548-TAG | 10 mM Glu + 100 uM BINA | 4 | 302 |
| Figure 4 | 548-TAG | 10 mM Glu + 5 uM Gi | 3 | 179 |
| Figure 5 | 359-463-TAG + HA-GSGS-mGluR3 | 10 uM Glu | 5 | 271 |
| Figure 5 | 359-463-TAG + HA-GSGS-mGluR4 | 10 uM Glu | 4 | 202 |
| Figure 5 | 359-463-TAG + HA-GSGS-mGluR7 | 10 uM Glu | 5 | 152 |
| Figure 5 | 359-463-TAG + HA-GSGS-mGluR4 | 1 mM LSP4-2022 | 4 | 197 |
| Figure 5 | 359-463-TAG + HA-GSGS-mGluR7 | 1 mM LSP4-2022 | 5 | 268 |
| Figure S3 | 359-463-TAG R177A + HA-GSGS-mGluR2-R177A | 1 uM Glu | 5 | 343 |
| Figure S3 | 359-463-TAG R177A + HA-GSGS-mGluR2-R177A | 10 uM Glu | 5 | 382 |
| Figure S3 | 359-463-TAG R177A + HA-GSGS-mGluR2-R177A | 100 uM Glu | 5 | 300 |
| Figure S3 | 359-463-TAG R177A + HA-GSGS-mGluR2-R177A | 1 mM Glu | 5 | 235 |
| Figure S3 | 359-463-TAG D95A + HA-GSGS-mGluR2-D95A | 0 mM Glu | 5 | 146 |
| Figure S3 | 359-463-TAG D95A + HA-GSGS-mGluR2-D95A | 1 uM Glu | 5 | 166 |
| Figure S3 | 359-463-TAG D95A + HA-GSGS-mGluR2-D95A | 10 uM Glu | 5 | 91 |
| Figure S3 | 359-463-TAG D95A + HA-GSGS-mGluR2-D95A | 100 uM Glu | 6 | 137 |
| Figure S3 | 359-463-TAG D95A + HA-GSGS-mGluR2-D95A | 1 mM Glu | 5 | 148 |
| Figure S3 | 359-463-TAG D95A + HA-GSGS-mGluR2-D95A | 10 mM Glu | 5 | 133 |
| Figure S3 | 359-463-TAG D95A + HA-GSGS-mGluR2-D95A | 20 uM LY37 | 5 | 70 |
| Figure S3 | 359-463-TAG + HA-GSGS-mGluR2 | 20 uM LY37 | 5 | 101 |

